# Metabolic commitment to human motor neurons is characterized by mitochondrial depletion and pyruvate rewiring

**DOI:** 10.64898/2026.10.05.756667

**Authors:** Ruben Torregrosa-Muñumer, Jeremi Turkia, Danil Spirin, Jouni Kvist, Jonna Saarimäki-Vire, Suleva Povea-Cabello, Rumeysa Ermis, Jana Pennonen, Emilia Kuuluvainen, Timo Otonkoski, Ville Hietakangas, Thomas G. McWilliams, Erika Fernandez-Vizarra, Pekka Katajisto, Henna Tyynismaa

## Abstract

Mitochondrial remodelling is a key feature of neurogenesis in cortical and hippocampal neuronal lineages, yet the temporal control of organellar homeostasis in developing human motor neurons remains poorly understood. Here, we generated a high-resolution temporal resource integrating proteomics, metabolomics, and mitochondrial characterization across human iPSC-to-motor neuron differentiation. This analysis revealed prolonged mitochondrial depletion during early neurogenesis, characterized by reduced mitochondrial content, mitochondrial fragmentation, and lysosome-dependent turnover. This state persisted throughout the neural progenitor stage and was followed by recovery of mitochondrial content during neuronal maturation. During this transition, pyruvate utilization shifted from pyruvate carboxylation towards pyruvate dehydrogenase-dependent oxidation. Pharmacological manipulation of pyruvate partitioning demonstrated that this metabolic switch is required for human neuronal differentiation. Our findings establish a temporal framework for the metabolic control of human motor neuron differentiation and reveal a progenitor-specific state linking mitochondrial depletion to pyruvate metabolism.

## INTRODUCTION

Mitochondria orchestrate stem cell fate by integrating bioenergetic, biosynthetic and signalling functions to coordinate developmental maturation (Arnold et al., 2022, Chandel et al., 2016, Dohla et al., 2022, Moussaieff et al., 2015, TeSlaa et al., 2016). Given their capacity for self-renewal and differentiation into all somatic lineages, pluripotent stem cells (PSC) are a powerful model to investigate how mitochondrial and metabolic programs are remodelled during cell fate transitions. PSC are classically depicted as harbouring immature mitochondria with limited oxidative capacity and a predominantly glycolytic phenotype, whereas differentiation triggers mitochondrial biogenesis, structural maturation and the shift to oxidative phosphorylation (OxPhos) (Cho et al., 2006, Facucho-Oliveira et al., 2007, Kelly et al., 2013, Prigione et al., 2010, St John et al., 2005, Varum et al., 2011, Zhang et al., 2011). However, accumulating evidence indicates that mitochondrial remodeling during lineage commitment is more complex than a universal transition from glycolytic pluripotent cells to oxidative differentiated cells (Birket et al., 2011, Kelly et al., 2013, Lees et al., 2018, Lv et al., 2022, Mostafavi et al., 2021). For example, mesodermal and endodermal differentiations exhibit increased OxPhos despite reduced mitochondrial abundance (Lv et al., 2022, Mostafavi et al., 2021). In contrast, early neurogenesis relies on sustained glycolysis (Birket et al., 2011, Lees et al., 2018), and the switch to mitochondrial metabolism and OxPhos only occurs later during neuronal maturation (Cliff et al., 2017, Lange et al., 2016, Rumpf et al., 2023, Scandella et al., 2023). Impaired mitochondrial metabolism also alters neurodevelopmental outcomes, demonstrating that metabolic programs actively regulate cell fate decisions rather than simply adapting to changing energetic demands (Beckervordersandforth, 2017, Homem et al., 2014, Iwata et al., 2020, Khacho et al., 2016, Petrelli et al., 2023, Stoll et al., 2015, Xu et al., 2023). Despite these insights, the temporal coordination of mitochondrial and metabolic remodelling during early neurogenesis remains incompletely understood. Specifically, how human motor neurons acquire the metabolic capacity required for their specialized physiology remains unclear.

Human induced pluripotent stem cell-derived motor neurons (iPSC-MN) provide a tractable system to study metabolic remodelling across discrete stages of developmental lineage progression (Amoros et al., 2022, Castillo Bautista and Sterneckert, 2022). Temporal multi-omics studies in NGN2-induced glutamatergic cortical-like neurons (iNeurons or iNs) have defined key metabolic changes during neuronal differentiation (Danacikova et al., 2025, Ordureau et al., 2021). Yet comparable stage-resolved analyses across directed motor neuron differentiation remain scarce.

Here, we profiled human motor neuron differentiation using high-resolution multi-omics. Our findings identify a previously unrecognized metabolic intermediate stage during human motor neuron differentiation, defined by transient mitochondrial depletion, fragmentation, autophagic remodelling, preferential pyruvate carboxylation, and delayed oxidative maturation. Rather than progressing gradually, metabolic maturation in motor neurons proceeds through a distinct developmental state where pyruvate utilization is actively reprogrammed. Crucially, promoting pyruvate dehydrogenase (PDH)-dependent pyruvate oxidation accelerated neuronal differentiation, establishing pyruvate metabolism as an active regulator of developmental progression. Moreover, this framework provides a comprehensive reference for investigating mitochondrial and metabolic dynamics during human neurogenesis and offers a foundation for future studies examining how mitochondrial signalling pathways regulate stem cell fate decisions. Together, these findings reveal a previously unexpected link between mitochondrial network remodelling and human motor neuron fate.

## RESULTS

### Transient mitochondrial depletion is a shared feature of early lineage commitment

Mitochondrial networks undergo extensive remodelling during differentiation and content typically increases as stem cells acquire mature cell identity. To define the temporal regulation of metabolic homeostasis in human motor neurons, we used a robust directed differentiation protocol that generates functional, electrically active iPSC-MN within 30 days (Guo et al., 2017, Harjuhaahto et al., 2020, Maury et al., 2015, Sainio et al., 2021). Sequential modulation of developmental pathways (Figure 1A) yielded motor neurons expressing ISL1/2 and HB9, confirming their spinal motor neuron identity (Figure 1B).

**Figure 1:**
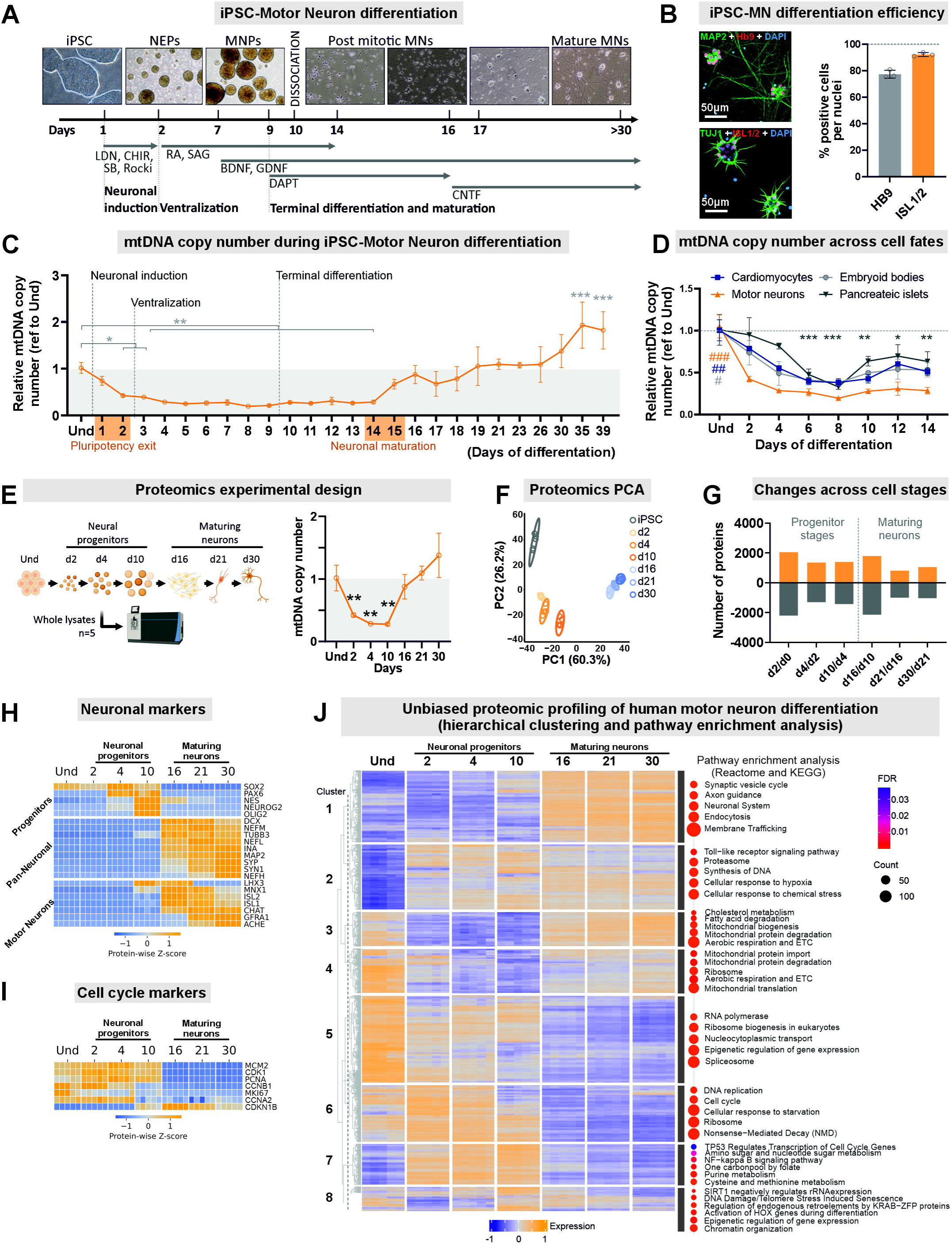
Mitochondrial content is reduced upon pluripotent stem cell differentiation. **A)** Cartoon depicting the directed differentiation protocol used to generate iPSC-MN. **B)** Validation of motor neuron differentiation by immunocytochemistry for the motor neuron markers ISL1/2 and Hb9 and the pan neural markers TUJ1 and MAP2. Left, representative images of iPSC-MN neurons at day 30 (iPSC 1 cell line). Right, quantification of Hb9 or ISL1/2 positive cells normalized to total nuclei from three preparations (n = 3 technical replicates), including a total of 24-29 images and 2240-3186 cells for Hb9 and ISL1/2 respectively. Cell line: iPSC1. **C)** mtDNA copy number was quantified by qPCR over 39 days of iPSC-MN differentiation and normalized to the undifferentiated starting point. Dotted lines indicate key developmental stages in the differentiation protocol. Cell line: iPSC 1; n = 3–6 technical replicates. **D)** mtDNA copy number was quantified by qPCR during the first 14 days of differentiation toward ectoderm (motor neurons, same data as in Figure 1C), mesoderm (cardiomyocytes), endoderm (beta cells), and during spontaneous differentiation in embryoid bodies. Each differentiation condition was normalized and compared to its corresponding undifferentiated control. n = 3-6 technical replicates. Cell lines: iPSC1 cell line was used in all differentiation, except for beta cells, where ESC H1 was used. **E)** On the left, proteomics experimental design showing the selected time points and number of technical replicates: five wells of the iPSC1 cell line per time point. Created in BioRender. Torregrosa, R. (2026) https://BioRender.com/qwqb483. On the right, mtDNA copy number values for the selected time points (same data as in Figure 1C). **F)** Principal component analysis (PCA) of the proteomics data showing clear separation between differentiation stages. **G)** Number of significantly changed proteins between consecutive transitions, with the largest changes observed at the d0 to d2 and d10 to d16 transitions. **H)** Heatmap from proteomics data showing selected markers of neuronal progenitors, pan neuronal markers, and motor neuron markers displayed as Z-scores. **I)** Heatmap from proteomics data showing selected cell cycle markers displayed as Z-scores. **J)** On the left, hierarchical clustering of expression trajectories for all detected proteins during motor neuron differentiation, with expression values transformed from 1 to −1. On the right, pathway enrichment analysis of the resulting clusters based on KEGG and Reactome databases. For all, error bars represent standard deviation. Statistical significance was defined as p < 0.05 (*), p < 0.01 (**), or p < 0.001 (***); # denotes significantly different compared to all groups. Multiple group comparisons were performed using one way ANOVA followed by Dunnett’s multiple comparison test (vs Und).

We performed a high-resolution temporal analysis of mtDNA copy number across 39 days of differentiation, which revealed a biphasic pattern of mtDNA remodelling (Figure 1C). First, mtDNA copy number declined dramatically during ectodermal specification (days 1–6). It then remained approximately threefold lower than in iPSC throughout the progenitor stage (days 6–14), regardless of whether cells were cultured in suspension (days 6–10) or as a monolayer (days 10–14). This observation refines the prevailing model of progressive mitochondrial expansion during stem cell differentiation. Notably, motor neuron progenitor commitment is accompanied by a substantial and prolonged reduction in mitochondrial content. Second, mtDNA copy number began to recover around days 14–15, coinciding with the onset of neuronal maturation (Figure 1C). We observed the same dynamics across independent iPSC and embryonic stem cells (ESC) lines (Supplementary Fig. 1A–B). A similar decrease in mtDNA copy number was previously reported during the differentiation of human pluripotent stem cells into neural precursor cells by Lees et al., who observed a transient reduction in mtDNA abundance and mitochondrial activity during early neural induction (Lees et al., 2018). However, the magnitude and duration of mtDNA depletion observed in our study were considerably larger, with mtDNA copy number remaining suppressed throughout the neural progenitor stage before recovering during motor neuron maturation. Moreover, omission of individual small molecules included in the differentiation protocol, either alone or in combination, did not prevent the changes in mtDNA copy number, indicating that these dynamics were not driven by any single differentiation cue (Supplementary Fig. 1C–F).

To determine whether the early decline in mtDNA was specific to neuroectodermal differentiation, we extended our analysis to cardiomyocytes, pancreatic islets, and embryoid bodies (EBs). Unlike directed differentiation protocols, EBs undergo spontaneous and asynchronous multi-lineage differentiation in suspension. Surprisingly, all models exhibited mtDNA depletion during early differentiation (Figure 1D). A similar decline was also observed *in vivo* during spontaneous differentiation in a teratoma assay (Supplementary Fig. 1G). Together, these findings indicate that early mtDNA depletion occurs across diverse differentiation systems and germ-layer trajectories, suggesting that it is a general feature of early lineage commitment rather than a neuroectoderm- or protocol-specific response.

To contextualise mtDNA dynamics across motor neuron differentiation, we profiled the proteome at representative developmental stages (Figure 1E), revealing stage-specific proteomic states (Figure 1F). The most extensive changes occurred between day 0 / 2, corresponding to exit from pluripotency, and between days 10 / 16, coinciding with motor neuron specification and the onset of neuronal maturation (Figure 1G). These transitions corresponded closely to the initial decline and subsequent recovery of mtDNA copy number (Figure 1C). Stage-specific markers confirmed progression from pluripotency to mature motor neurons (Figure 1H). Cell-cycle markers showed that early progenitors were highly proliferative, whereas neurons became post-mitotic after day 10 (Figure 1I).

Hierarchical clustering of temporal protein-expression trajectories, followed by KEGG and Reactome pathway enrichment analysis, identified eight distinct clusters (Figure 1J and Supplementary Figure 1H). Cluster 1 progressively increased during differentiation and was enriched for neurogenesis-related pathways. Cluster 2 increased sharply following pluripotency exit and included proteins involved in proteasomal function and responses to hypoxia and chemical stress. Clusters 3 and 4 were enriched for mitochondrial pathways and followed trajectories that broadly paralleled mtDNA copy number, supporting coordinated remodelling of mtDNA and mitochondrial protein abundance. Cluster 5 declined progressively and was enriched for RNA processing and epigenetic regulation. Cluster 6 contained proteins abundant in iPSC and early progenitors and was enriched for DNA replication, translation, and cell-cycle regulation. Cluster 7 followed an inverse trajectory to mitochondrial abundance and was enriched for amino acid, purine, and one-carbon metabolism, consistent with the biosynthetic demands of proliferating progenitors. Finally, Cluster 8 peaked around day 10 and was enriched for chromatin-remodelling pathways.

PSC are often described as having lower mitochondrial content and reduced OxPhos capacity in comparison to parental fibroblasts (Armstrong et al., 2010, Kelly et al., 2013, Prigione et al., 2010, Prigione et al., 2011). In contrast, we observed higher mtDNA levels in iPSC than in parental fibroblasts, and similar levels between ESC and iPSC, consistent with (Birket et al., 2011), regardless of the culture conditions (Supplementary Fig. 2A-D). Mitochondrial protein levels were likewise higher in iPSC and ESC compared to parental fibroblasts (Supplementary Fig. 2E-F). Nevertheless, stem cells remained predominantly glycolytic and displayed immature mitochondrial morphology and function, consistent with previous reports. (Armstrong et al., 2010, Kelly et al., 2013, Prigione et al., 2010, Prigione et al., 2011) (Supplementary Fig. 2G-P). Yet, despite their immature phenotype, mitochondria in PSC remained metabolically active (Supplementary Fig. 2J and (Zhang et al., 2011)). Thus, mitochondrial abundance alone does not define the metabolic state of PSC.

Together, the mtDNA and proteomic analyses revealed two major phases of mitochondrial remodelling during motor neuron differentiation: an early reduction associated with pluripotency exit and progenitor formation, followed by mitochondrial recovery during neuronal specification and maturation. More broadly, this temporal dataset provides a framework for investigating the molecular changes accompanying human neuronal differentiation and mitochondrial remodelling.

### The metabolic transition state is accompanied by mitochondrial fragmentation and autophagic remodeling

Next, we confirmed key proteomic observations by immunoblotting in independent iPSC lines. The levels of the pluripotency marker Nanog were rapidly reduced upon differentiation induction, while SOX1 (an early ectoderm specification marker) showed a progressive increase, confirming lineage commitment (Figure 2A). Similar to the reduction in mtDNA content, mitochondrial mass decreased during differentiation (Figure 2A, B). Selected OxPhos subunits followed a similar pattern, although the decline was less pronounced for some subunits (Figure 2A, C).

**Figure 2:**
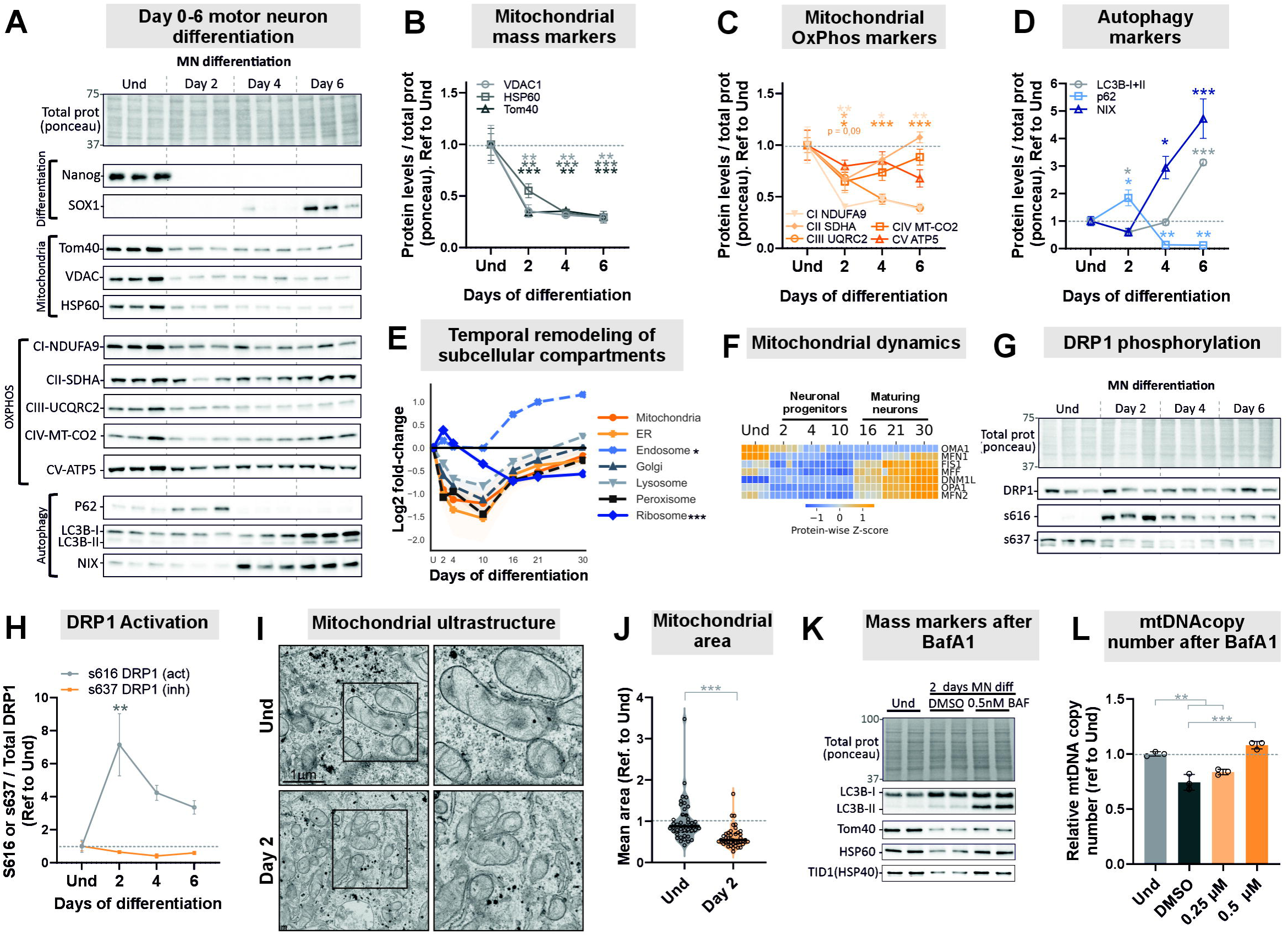
The metabolic transition state is accompanied by mitochondrial fragmentation and autophagic remodeling. **(A)** Western blot analysis of iPSC-MN during the first six days of differentiation (days 0, 2, 4, and 6). n = 3 biological replicates (iPSC1–3). Total protein normalization was performed using Ponceau staining. Differentiation status was assessed by Nanog (pluripotency marker) and SOX1 (early neuroectoderm marker). Multiple mitochondrial mass markers were analyzed, including outer mitochondrial membrane proteins (TOM40 and TOM20) and the mitochondrial matrix protein HSP60. In addition, one representative subunit from each OxPhos complex was examined (CI = Complex I, etc.). Autophagy-related proteins analyzed included p62, LC3B and its lipidated form (LC3-II), and NIX (BNIP3L). **(B)** Quantification of mitochondrial mass markers levels from panel (A), normalized to total protein (Ponceau). Each marker was further referred to and compared with its corresponding undifferentiated (Und) levels. **(C)** Quantification of OxPhos subunits levels from panel (A), normalized to total protein (Ponceau). Each marker was further referred to its corresponding undifferentiated (Und) levels **(D)** Quantification of autophagy-related markers from panel (A), normalized to total protein and expressed relative to their corresponding Und levels. LC3B abundance was calculated as the sum of LC3-I and LC3-II. Each marker was normalized to total protein (Ponceau) and further referred to its corresponding undifferentiated (Und) levels. **(E)** Temporal remodelling of subcellular compartments during differentiation from the proteomics data. Proteins were assigned to subcellular modules using previously reported compartment annotations (Ordureau et al., 2021), with related compartments collapsed to reduce redundancy (mitochondrial inner membrane, matrix, and outer membrane → *Mitochondria*; ER lumen, membrane, and high-curvature → *ER*; ER–Golgi intermediate compartment and cis-Golgi → *Golgi*). Protein abundances were log□ -transformed and normalized to the mean abundance of each protein across the five day-0 samples. Normalized protein values were subsequently averaged within each organelle category for each well and time point (*n* = 5 wells per time point). For each organelle category, paired differences relative to mitochondria were calculated within each sample. These differences were modelled using OLS regression with day as a categorical variable and day 0 as the reference level. HC3 robust standard errors were used for inference. Overall temporal changes were assessed using joint F-tests of the six post-baseline day coefficients, followed by Benjamini–Hochberg correction across organelle comparisons. Shaded areas represent SD. **(F)** Heatmap of proteomics data showing individual mitochondrial dynamics–related factors, displayed as Z-scores. **(G)** Western blot analysis performed on freshly extracted samples from iPSC1-3 as in panel (A). Mitochondrial fission regulator DRP1 was evaluated by analyzing total DRP1 levels as well as activating (Ser616) and inhibitory (Ser637) phosphorylation, using total protein (Ponceau staining) as loading control. **(H)** Quantification of DRP1 from panel (D), performed as described in (E). (**I)** Representative transmission electron microscopy images of undifferentiated iPSC1 (day 0) and day 2 differentiated neurons cultured as a monolayer, including magnified views. Scale bar = 1 µm. **(J)** Mitochondrial mean area between undifferentiated iPSC (day 0) and day 2 neural progenitors cultured as a monolayer, extracted from (I). n = 42 images containing 439 (day 0) or 272 (day 2) mitochondria from iPSC1 derived neurons. Differences between time points were assessed using an unpaired Welch’s t-test on log-transformed image-level means. **(K)** Western blot analysis was performed to assess lysosomal acidification following treatment with Bafilomycin A1 (BafA1) during the first 2 days of MN differentiation of iPSC1. Data represent *n* = 2 technical replicates (wells) from iPSC1. Inhibition of lysosomal acidification with 0.5 nM BafA1 resulted in the expected accumulation of lipidated LC3B (LC3B-II), confirming effective lysosomal blockade. Consequently, degradation of mitochondrial proteins was partly prevented, as indicated by elevated levels of the outer-membrane marker TOM40, and the matrix-localized proteins HSP60 and HSP40. **(L)** mtDNA copy number determined by qPCR in iPSC-derived neurons after 2 days of BafA1 treatment. Cell line: iPSC1; n = 3 technical replicates (wells). For all, error bars represent standard deviation. Statistical significance was defined as p < 0.05 (*), p < 0.01 (**), or p < 0.001 (***); # denotes significantly different compared to all groups. Multiple group comparisons were performed using one way ANOVA followed by Dunnett’s multiple comparison test unless otherwise indicated.

We then examined the mechanisms underlying mitochondrial content reduction. If mitochondrial biogenesis fails to keep pace with cell division, mitochondrial content would be progressively diluted. However, dilution alone could not explain the observed reduction, as mtDNA increased at a similar rate to nuclear DNA until day 2, despite a ∼50% reduction in mtDNA copy number (Figure 1C) and mitochondrial mass (Figure 2B) and only slowed thereafter (Supplementary Fig.3A).

Alternatively, selective mitochondrial degradation through mitophagy may contribute to the reduction in mitochondrial content (Martens and Ganley, 2026). Mitophagy is increasingly recognized as a key mechanism of metabolic remodelling during cell fate transitions. Indeed, programmed mitophagy has been reported in several differentiation models, including cardiomyocytes, myoblasts, and retinal ganglion cells (Esteban-Martinez et al., 2017, Lampert et al., 2019, Sin et al., 2016, Yazdankhah et al., 2021, Zhao et al., 2020). Our proteomic analysis detected multiple autophagy-related proteins (Supplementary Fig. 3B) and revealed a modest enrichment of an autophagy related pathway at the onset of differentiation (Cluster 3, Figure 1J; Reactome R HSA 9612973, Autophagy, q value = 0.0008). Several proteins involved in autophagy initiation, cargo recognition, and lysosomal function (e.g., ULK1, SQSTM1, and LAMP1) peaked at day 2, whereas proteins associated with autophagosome formation and expansion (e.g., BECLIN1, ATG7, and ATG12) increased progressively until day 10. Together, these findings support a role for autophagy during early neurogenesis.

To investigate whether mitochondrial turnover contributes to mitochondrial remodelling, we further measured autophagy and mitophagy markers by immunoblotting, including proteins not detected by proteomics. While p62 showed a transient peak at day 2, LC3B and the mitophagy receptor NIX (BNIP3L) displayed a progressive accumulation during the first 6 days of differentiation (Figure 2A and D). As NIX is a key regulator of programmed mitophagy during metabolic remodelling (Esteban-Martinez et al., 2017, Lampert et al., 2019, Schweers et al., 2007, Sin et al., 2016, Yazdankhah et al., 2021, Zhao et al., 2020), its sustained upregulation suggests that mitochondrial turnover pathways may be engaged during early differentiation. Interestingly, the proteomes of several organelles, including the endoplasmic reticulum, Golgi apparatus, lysosomes, and peroxisomes, followed temporal trajectories similar to those of mitochondria. In contrast, endosomal and ribosomal proteins displayed distinct patterns (Figure 2E).

Mitochondrial fragmentation often precedes mitophagy and is largely regulated by the mitochondrial fission protein DRP1 (Burman et al., 2017, Twig et al., 2008). Most regulators of mitochondrial dynamics displayed a pattern similar to mitochondrial mass markers (Figure 2F). However, DRP1 (DNM1L) was not reduced at the onset of differentiation. Instead, phosphorylation at Ser616 increased, consistent with DRP1 activation (Figure 2G,H). This was accompanied by a reduction in mitochondrial area, length, and width upon exit from pluripotency (Figure 2I,J and Supplementary Fig. 3C). Mitochondrial ultrastructure continued to change during the progenitor phase. Mitochondrial area reached a minimum at day 4 and remained low until day 10 (Supplementary Fig. 3F-J). These findings indicate that mitochondrial fragmentation accompanies the reduction in mitochondrial content and may facilitate organelle turnover through autophagic pathways.

We then asked whether autophagy contributes to the depletion of mitochondrial content during early differentiation. To inhibit autophagic flux, we blocked lysosomal V-ATPase activity using bafilomycin A1 (BafA1) (Yoshimori et al., 1991). BafA1treatment caused the accumulation of lipidated LC3B (LC3B-II) (Figure 2K), confirming effective inhibition of lysosomal degradation. Importantly, BafA1 markedly attenuated the decline in mitochondrial mass during the first 48 to 96 hours of differentiation (Figure 2L and Supplementary Fig. 3K-L). These findings demonstrate that lysosome-dependent turnover contributes substantially to early mitochondrial remodelling.

Given the induction of NIX during early differentiation and its established role in mitochondrial turnover, we generated NIX knockout and NIX/BNIP3 double-knockout iPSC and differentiated them into neurons. BNIP3 was included because it is a closely related mitophagy receptor with partially overlapping functions (Li et al., 2021, Uoselis et al., 2023). Neither individual loss of NIX nor the combined loss of NIX and BNIP3 altered mtDNA copy-number dynamics or differentiation efficiency in motor neurons (Supplementary Fig. 3M–R). Our findings suggest NIX and BNIP3 are not required for early mitochondrial depletion or lineage commitment in human motor neurons.

In sum, these findings link early mitochondrial depletion to DRP1-associated fragmentation and lysosome-dependent turnover. BafA1 largely prevented the loss of mitochondrial mass and mtDNA, whereas deletion of NIX alone or together with BNIP3 had no detectable effect. Thus, early mitochondrial remodelling in motor neurons depends on lysosomal activity but does not require NIX or BNIP3 under these conditions. Alongside the broad temporal remodelling of other organelle compartments in motor neurons, a defined window of non-selective macroautophagy may also account for this phenotype.

### The early transition state is characterized by suppression of oxidative pyruvate metabolism

Using our proteomics dataset and MitoCarta 3.0 annotations (Rath et al., 2021), we next examined how the OxPhos system is remodelled during neural differentiation. The OxPhos system comprises the electron transport chain complexes I–IV and ATP synthase (complex V), each consisting of structural subunits and dedicated assembly factors. Consistent with the mitochondrial content dynamics (Figure 1C and J), virtually all OxPhos subunits were enriched in iPSC, decreased at the progenitor stage, and progressively increased during neuronal maturation (Figure 3A). Assembly factors broadly followed the same trend but with greater variability, suggesting complex-specific patterns (Figure 3B). Comparison of subunit and assembly factor trajectories revealed distinct regulatory patterns across OxPhos complexes. Complexes I and III showed strong concordance throughout differentiation, whereas Complexes IV and V diverged slightly during the progenitor phase, with assembly factors remaining relatively preserved (Figure 3C). In contrast, complex II displayed minimal correlation, with the assembly factors (SDHAF1, SDHAF2 and SDHAF4) increasing early during differentiation (Figure 3B). Notably, mtDNA-encoded proteins showed a stronger transient reduction during early differentiation than nuclear-encoded mitochondrial proteins. As these proteins form the catalytic core of the respiratory complexes, this pattern suggests selective remodelling of the OxPhos proteome during neuronal commitment. (Figure 3D).

**Figure 3:**
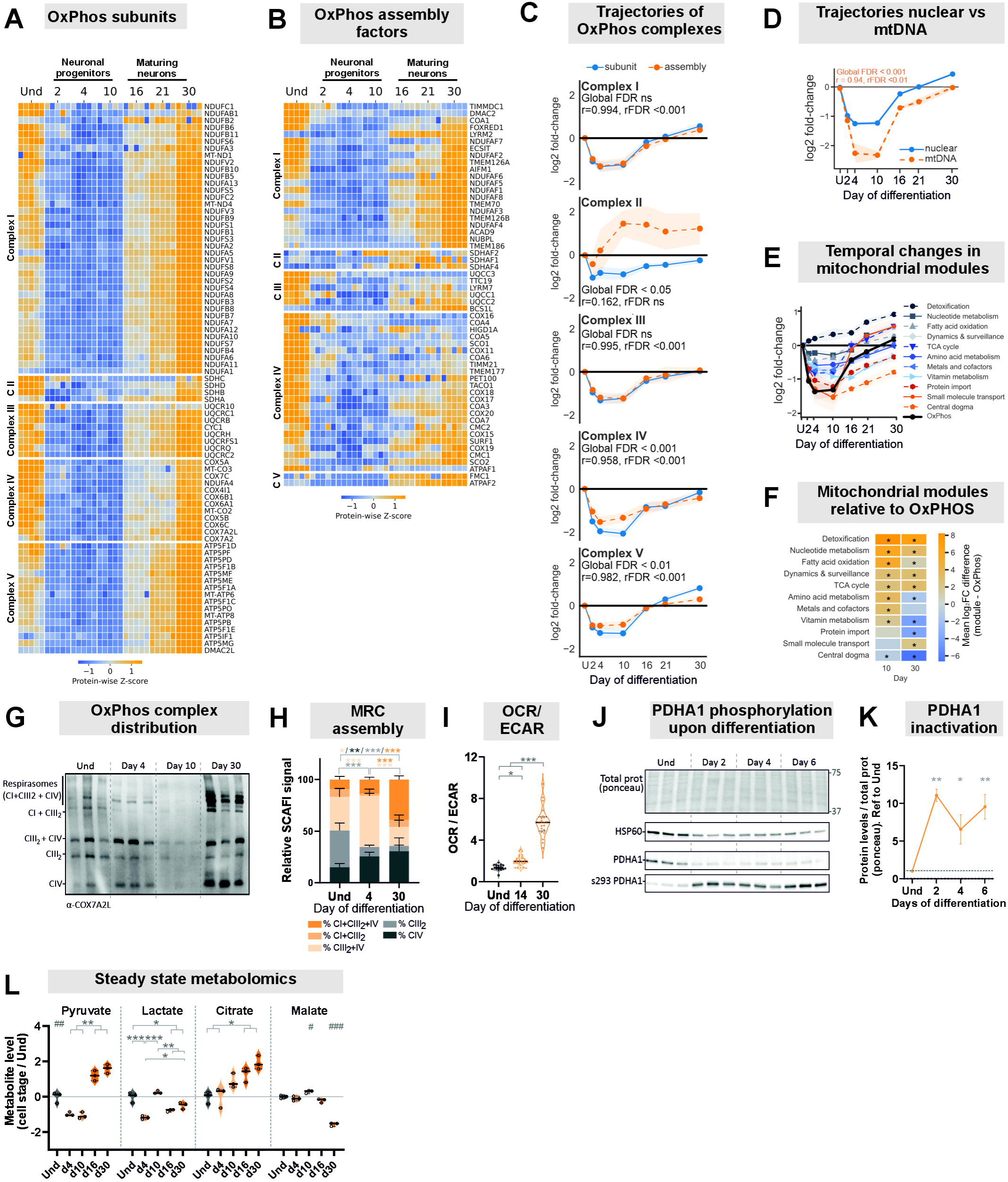
Coordinated remodeling of mitochondrial OxPhos complexes during neuronal maturation. **(A–B)** Heatmaps of proteomics data showing OxPhos subunits **(A)** and OxPhos assembly factors **(B)**, grouped by respiratory chain complex (Complex I–V) according to MitoCarta 3.0 annotations. Protein abundances were scaled and displayed as Z-scores across differentiation time points to highlight coordinated temporal changes within and across complexes. **(C)** Trajectories of OxPhos subunits and assembly factors for each complex using MitoCarta 3.0 annotations. Protein abundances were log□-transformed and normalized to the mean abundance of each protein across the five day-0 samples. Protein values were subsequently averaged within each OxPhos complex and protein category for each well and time point (n = 5 wells per time point). Differences between assembly factors and subunits were calculated within each sample. Temporal changes in these paired differences were analysed separately for each complex using a OLS regression model with day as a categorical variable and day 0 as the reference level. HC3 robust standard errors were used for inference. Overall temporal changes were assessed using joint F-tests of the six post-baseline day coefficients, followed by Benjamini–Hochberg correction across organelle comparisons.. Shaded areas represent SD. **(D)** Trajectories of nuclear- and mtDNA-encoded OxPhos subunits using MitoCarta 3.0 annotations. Data processed as in (C). **(E)** Mitochondrial pathways trajectories across differentiation. Data processed as in (C). Mitochondrial proteins were annotated using MitoCarta 3.0 MitoPathways and collapsed into broader functional modules for temporal analysis. Proteins annotated to OXPHOS or electron carrier pathways were classified as OxPhos; mitochondrial central dogma, mitochondrial ribosome, mitochondrial translation, and RNA-processing pathways as Central dogma; TCA cycle, citric acid cycle, Krebs cycle, and carbohydrate metabolism as TCA cycle; fatty acid oxidation, β-oxidation, FAO, and lipid metabolism as Fatty acid oxidation; and mitochondrial dynamics, dynamics and surveillance, mitophagy, fission, fusion, PINK1, and PRKN annotations as Dynamics & surveillance. Protein import, TOM/TIM complex, and protein-homeostasis annotations were grouped as Protein import. Amino acid metabolism, nucleotide metabolism, metals and cofactors, vitamin metabolism, detoxification, small-molecule transport, and their corresponding MitoCarta annotations were retained as individual modules. **(F)** Relative remodelling of mitochondrial modules at days 10 and 30. For each well, the mean OxPhos log□ fold change was subtracted from the mean log□ fold change of each mitochondrial module. Heatmap values represent the mean paired difference across five independent wells. Positive values indicate a greater increase or smaller reduction relative to OxPhos, whereas negative values indicate a greater reduction or smaller increase. Differences were assessed using paired two-sided *t*-tests, with Benjamini–Hochberg correction across all comparisons within each time point. Asterisks indicate p<0.01. **(G)** Representative Blue-Native PAGE (BN-PAGE) immunoblot showing respiratory chain complexes organization. SCAFI / COX72AL is shown as a marker of Complex III and IV–containing assemblies. n = 3 biological replicates (iPSC1-3). **(H)** Mitochondrial respiratory chain (MRC) assembly. Quantification of the BN-PAGE analysis shown in (G), depicting the relative abundance (%) of respirasomes (CI+CIII□ +CIV), supercomplexes I+III_2_ and III_2_+IV, and individual respiratory chain complexes. Quantification of day 10 samples was excluded because low mitochondrial abundance resulted in poor BN-PAGE resolution. **(I)** The oxygen consumption rate (OCR)–to–extracellular acidification rate (ECAR) ratio was calculated from Seahorse XF measurements to assess metabolic preference between OxPhos and glycolysis, with lower ratios indicating a more glycolytic metabolic phenotype. Only differentiation stages growing as monolayers were analyzed to ensure comparable measurement conditions. Data represent *n* = 18–30 wells pooled from two independent cell lines (iPSC1-2). **(J)** Western blot analysis of iPSC-MN during the first six days of differentiation (days 0, 2, 4, and 6). n = 3 biological replicates (iPSC1–3). HSP60 was used to confirm mitochondrial content reduction during differentiation. Activity of the mitochondrial PDH complex was assessed by measuring total PDHA1 subunit levels and phosphorylation at Ser293 (inactivating). **(K)** Quantification of PDHA1 from panel (E). Signals for each antibody were first normalized to total protein (Ponceau staining). The ratio of phosphorylated PDHA1 to total PDHA1 was then calculated and normalized to undifferentiated (Und) levels. **(L)** Intracellular abundance of glycolysis (pyruvate and lactate) and TCA cycle metabolites (citrate and malate), shown as log2-transformed values normalized to undifferentiated (Und) cells. Cell line: iPSC1; n = 2-3 technical replicates (wells). Data are presented as mean ± standard deviation. Statistical significance was defined as p < 0.05 (*), p < 0.01 (**), or p < 0.001 (***). Multiple group comparisons were performed using ANOVA followed by Dunnett’s (F) or Tukeýs (H) multiple comparison test.

Although the mitochondrial proteome was broadly reduced during early differentiation, the magnitude of this reduction varied across mitochondrial proteins. We therefore examined how distinct mitochondrial pathways were remodelled during neuronal differentiation. Grouping mitochondrial proteins into functional pathways based on MitoCarta 3.0 revealed that most pathways underwent a transient decline during early progenitor stages followed by recovery during neuronal maturation (Figure 3E and Supplementary Figure 4A). However, the magnitude of suppression differed between pathways. Pathways linked to amino acid, nucleotide and fatty acid metabolism showed more modest reductions than OxPhos-related proteins. These findings indicate that mitochondrial remodelling during progenitor expansion is not uniform, with OxPhos components undergoing the most pronounced reduction. In contrast, the mitochondrial detoxification pathway progressively increased throughout differentiation, highlighting the activation of antioxidant and stress response programmes (Figure 3E and Supplementary Figure 4A).

To determine whether these metabolic changes were accompanied by remodelling of the respiratory chain, we examined the organization of OxPhos complexes by Blue-Native PAGE (Figure 3G–H and Supplementary Fig. 4 B-C). Early differentiation stages were characterized by a predominance of free CIII□, CIV and supercomplex (SC) III□+IV. At later neuronal maturation stages, free CIV, SC I+III_2_ and respirasomes (I+III□+IV) prevailed. Thus, the recovery of mitochondrial content during neuronal maturation was accompanied by increased Complex IV assembly and a relative enrichment of respiratory supercomplexes. Consistent with these structural changes, the OCR/ECAR (oxygen consumption rate / extracellular acidification rate) ratio increased progressively from day 14 to day 30 (Figure 3I), indicating a gradual shift towards a more oxidative metabolic state during neuronal maturation. These findings show that neuronal maturation is accompanied by coordinated remodelling of mitochondrial content, respiratory chain organization, and cellular metabolism. Whether the increased abundance of respiratory supercomplexes directly contributes to this oxidative phenotype or simply reflects the higher mitochondrial content of mature neurons remains unclear.

Because pyruvate links glycolysis to mitochondrial metabolism, we next examined how pyruvate entry into oxidative metabolism through the PDH complex is regulated upon pluripotency exit. PDHA1 protein levels transiently decreased during the progenitor stage, mirroring the reduction in OxPhos components. At the same time, inhibitory phosphorylation of PDHA1 at Ser293 increased significantly (Figure 3J,K), suggesting that pyruvate entry into mitochondrial oxidative metabolism is restricted during the progenitor stage.

To determine how these proteomic changes are reflected at the metabolite level, we performed steady-state targeted metabolomics across the differentiation time course (Figure 3L and Supplementary Fig. 4 D-G). Pathway-level analysis revealed broad metabolic remodelling during differentiation, with amino acid metabolism among the most prominently affected metabolic categories (Supplementary Fig. 4F). At the metabolite level, however, glycolytic and tricarboxylic acid (TCA) cycle intermediates displayed heterogeneous trajectories rather than coordinated changes. Citrate progressively accumulated during differentiation, whereas malate levels declined. Glycolytic intermediates also displayed distinct temporal profiles, with pyruvate decreasing early during differentiation and lactate showing a transient increase at intermediate stages. These findings reveal extensive metabolic remodelling during neuronal differentiation and suggest that changes in metabolite abundance alone are insufficient to infer pathway activity.

These data reveal that motor neuron differentiation is accompanied by at least two major phases of mitochondrial metabolic remodelling. Early progenitor commitment is characterized by transient suppression of mitochondrial oxidative metabolism and regulation of pyruvate utilization. In contrast, neuronal maturation is associated with progressive mitochondrial biogenesis, respirasome assembly, and increased oxidative capacity. These findings support a model in which mitochondrial metabolism is dynamically reprogrammed during neurogenesis to accommodate the distinct metabolic demands of progenitor expansion and neuronal maturation.

### A switch from pyruvate carboxylation to pyruvate oxidation accompanies neuronal maturation

Leveraging the insights from our proteomic dataset, we next asked how central carbon metabolism is remodelled during differentiation (Figure 4A). Early progenitor stages were characterized by increased expression of glucose transporters, primarily SLC2A1/GLUT1 and SLC2A4/GLUT4, together with a transient peak in the lactate exporter SLC16A3/MCT4 (Dimmer et al., 2000). This pattern is consistent with increased glycolytic activity during early differentiation (Cliff et al., 2017, Zheng et al., 2016). In contrast, neuronal maturation was associated with increased expression of the mitochondrial pyruvate importer MPC2, SLC2A6/GLUT6, and a moderate recovery of SLC2A3/GLUT3, suggesting enhanced mitochondrial pyruvate utilization. Consistent with these transporter changes, glycolytic enzymes and related branch pathways were most abundant during progenitor stages (Figure 4B and Supplementary Fig. 5A and B), whereas TCA cycle proteins progressively increased during neuronal maturation (Figure 4C). These findings are consistent with exit from pluripotency during ectoderm commitment being associated with a glycolytic and fermentative metabolic state (Cliff et al., 2017), whereas neuronal maturation is associated with progressively increased mitochondrial oxidative metabolism. Nevertheless, glycolysis remained a central pathway throughout differentiation, likely providing carbon substrates for mitochondrial and intermediates for redox and biosynthetic pathways. Consistent with this sustained but remodelled role of glycolysis, neuronal maturation was accompanied by glycolytic isoform switching. This included the previously described HK2-to-HK1 and ENO1-to-ENO2 transitions (Zheng et al., 2016), as well as an ALDOA-to-ALDOC transition (Figure 4B).

**Figure 4:**
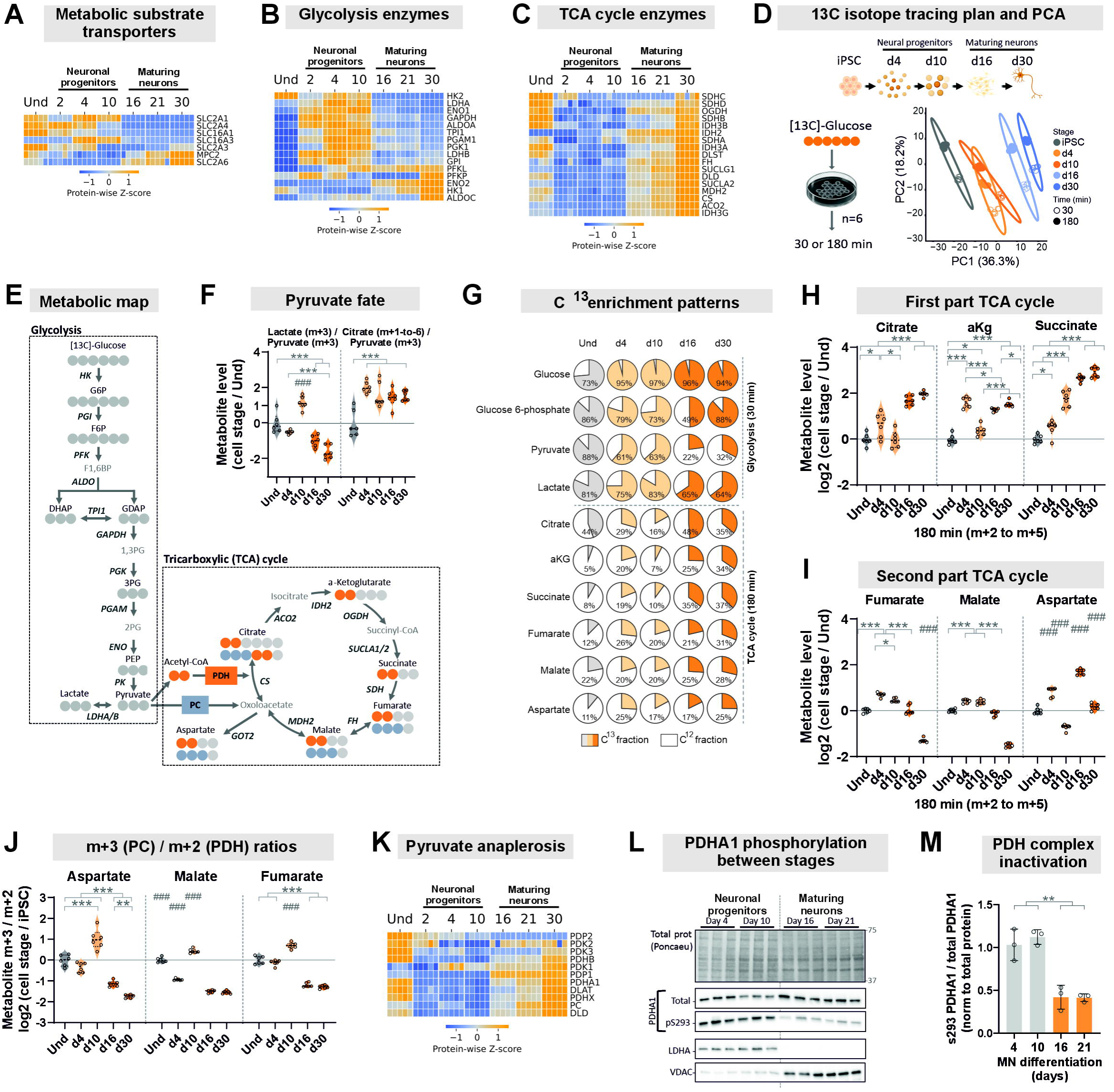
Metabolic shift from glycolysis-dominant early progenitors to enhanced OxPhos and TCA anaplerosis via PDH during neuronal maturation. **A)** Heatmap from proteomics data showing selected markers of substrate transports (SLCA16, glucose; SLC2A, pyruvate; and MPC2, lactate) displayed as Z-scores. **B)** Heatmap from proteomics data showing selected glycolysis enzymes displayed as Z-scores. **C)** Heatmap from proteomics data showing selected TCA cycle enzymes displayed as Z-scores. **D)** Experimental design for isotopic tracing metabolomics showing the selected time points and number of technical replicates: six wells of the iPSC1 cell line per time point. Cells were cultured for 30 or 180 minutes in medium containing 21.25 mM U^13^C-glucose. Created in BioRender. Torregrosa, R. (2026) https://BioRender.com/qwqb483. Lower left, principal component analysis (PCA) of the isotopic tracing metabolomics data showing clear separation between differentiation stages. **E)** Cartoon schematic of glycolysis and TCA cycle metabolites and enzymes. Shadowed circles denote carbon atoms. In the TCA cycle, orange circles indicate two ^13^C-labeled carbons derived from the PDH complex and acetyl-CoA, whereas blue circles indicate ^13^C labelled carbons originating from PC; metabolites without circles were not detected in our dataset. **F)** Intracellular metabolite ratios of m+3 lactate to m+2 pyruvate, or of labelled citrate (m+1 to m+6) to total citrate, used to infer pyruvate utilization; lower ratios indicate reduced pyruvate usage or preference. 180 minutes labelling. **G)** Pie charts of intracellular ^13^C enrichment patterns in glycolytic (30 minutes labelling) and TCA metabolites (180 minutes labelling) showing the total labelled ^13^C fraction (m+1 to m+n) in shaded colors (grey for undifferentiated cells, light orange for neuronal progenitors, and dark orange for maturing neurons), with the unlabelled ^12^C fraction in white; each pie represents 100% of the metabolite pool. The indicated % number reflects the total labelled fraction. **H)** Intracellular relative abundance of ^13^C-labeled carbons (m+2 to m+n, labelled fraction derived from pyruvate) after 180 minutes in TCA clockwise metabolites (citrate, α-ketoglutarate (αKg), and succinate), shown as log2-transformed values normalized to undifferentiated (Und) cells. **I)** Intracellular relative abundance of ^13^C-labeled carbons (m+2 to m+n, labelled fraction derived from pyruvate) after 180 minutes in TCA counterclockwise metabolites (fumarate, malate and aspartate (proxy of oxoloacetate), shown as log2-transformed values normalized to undifferentiated (Und) cells. **J)** Intracellular metabolite ratio m+3 to m+2 of aspartate, malate and fumarate. PC anaplerosis is counting for m+3, whereas PDH for m+2. A low ratio denotes PDH preference. **K)** Heatmap derived from proteomics data showing selected components of pyruvate anaplerosis, including PC (pyruvate carboxylase), the PDH complex (DLAT, DLD, PDHA1, PDHB, PDHX), and its regulatory enzymes, PDKs (inhibitory kinases) and PDPs (activating phosphatases). **L)** Western blot analysis of iPSC-MN across differentiation (days 4–21; *n* = 3 biological replicates, iPSC1–3). Total protein normalization was performed using Ponceau staining. Total PDHA1 and phosphorylated PDHA1 (pS293) were used to evaluate PDH activity, as phosphorylation at S293 inhibits the complex. LDHA and VDAC were used as markers of glycolysis and mitochondria, respectively. **M)** Quantification pS293/total PDHA from B. Data are presented as mean ± standard deviation. Statistical significance was defined as p < 0.05 (*), p < 0.01 (**), or p < 0.001 (***); # denotes significantly different compared to all groups. Multiple group comparisons were performed using one way ANOVA followed by Tukey’s multiple comparison test.

Although proteomic analyses revealed a robust upregulation of glycolytic enzymes during the progenitor stages, steady-state metabolomics did not show a corresponding accumulation of glycolytic intermediates (Figure 3L). This discrepancy suggests that metabolite abundance alone may not accurately reflect pathway activity. We therefore examine glucose carbon utilization using isotope tracing with uniformly ^13^C-labelled glucose across differentiation (Figure 4D-E and Supplementary Fig. 5C). To account for differences in labelling kinetics, glycolytic metabolites were analysed after 30 minutes of ^13^C-glucose labelling, whereas TCA cycle intermediates were analysed after 180 minutes. In iPSC and early progenitors, pyruvate-to-lactate conversion was higher than in more mature neurons (Figure 4F), and extracellular lactate accumulated more rapidly, particularly at day 10 (Supplementary Fig. 5D). These findings are consistent with a more fermentative glycolytic state during early differentiation. Despite the glycolytic phenotype of early progenitors, the citrate-to-pyruvate ratio increased rapidly upon differentiation (Figure 4F), implying that mitochondrial pyruvate metabolism is also engaged early during lineage commitment. Interestingly, this increase occurred despite the reduced abundance of OxPhos proteins at the onset of differentiation, suggesting that mitochondrial carbon metabolism is maintained independently of respiratory chain expansion. Consistent with this interpretation, components of the cytosol-mitochondria carbon transfer pathways, including ME1, GOT1 and MDH1, were transiently upregulated during the progenitor transition (Supplementary Fig. 5E–F).

Intracellular fractional enrichment analysis revealed stage-specific routing of glucose-derived carbon. Early progenitors directed a greater fraction of glucose-derived carbons into glycolysis, especially in pyruvate and lactate, whereas neuronal maturation was associated with increased incorporation into TCA cycle intermediates, especially citrate-succinate (Figure 4G), consistent with the corresponding enzyme levels (Figure 4B and C). Stable or reduced glycolytic metabolite abundance despite increased glucose labelling (Supplementary Fig. 5G) suggests rapid turnover and consumption. Within the TCA cycle, glucose-derived metabolites of the first half (citrate–succinate) progressively accumulated during maturation (Figure 4H), consistent with enhanced oxidative metabolism. In contrast, labelling of second-half intermediates (fumarate-malate) was highest in early progenitors (Figure 4I), indicating differential routing of glucose-derived carbon through the TCA cycle during differentiation.

Pyruvate is a central metabolic node whose mitochondrial metabolism can proceed through either PDH or pyruvate carboxylase (PC), thereby influencing carbon flow through the TCA cycle and related biosynthetic pathways (Arnold and Finley, 2023, Inigo et al., 2021). These alternative metabolic routes differentially support energy production, biosynthesis, and anaplerosis. Consistent with this, increasing evidence indicates that pyruvate metabolism contributes to stem cell state transitions and neuronal differentiation (Arnold et al., 2022, Iwata et al., 2023, Petrelli et al., 2023, Stoll et al., 2015, Zheng et al., 2016).

Because PDH- and PC-mediated pyruvate metabolism generate distinct m+2 and m+3 isotopologue signatures, respectively (Figure 4E), we quantified the intracellular m+3/m+2 ratio to assess the relative contribution of these pathways during neuronal differentiation (Figure 4J). Early progenitors displayed elevated intracellular m+3/m+2 ratios, indicative of preferential pyruvate carboxylation, with the ratio reaching its maximum at day 10. Thereafter, the m+3/m+2 ratio progressively declined during neuronal maturation, consistent with an increasing contribution of PDH-mediated pyruvate oxidation. Confirming this, proteins involved in PDH-mediated pyruvate oxidation were enriched during neuronal maturation (Figure 4K), while inhibitory phosphorylation of PDHA1 at Ser293 progressively decreased (Figure 4L-M), indicating increased PDH activity at later stages. Collectively, these findings suggest that early progenitors preferentially partition glucose-derived carbon towards PC, whereas mature neurons increasingly utilize pyruvate through PDH.

The early increase in α-ketoglutarate (αKG) at day 4 (Figure 4H) was particularly intriguing, as αKG is an essential co-substrate for DNA and histone demethylases, whereas succinate acts as a competitive inhibitor of these enzymes (Baksh and Finley, 2021, TeSlaa et al., 2016). Consistent with this possibility, αKG-to-succinate ratio was significantly elevated at day 4 (Supplementary Fig. 5H), suggesting a metabolic environment favourable for αKG-dependent dioxygenase activity. Supporting this posibility and the findings of (TeSlaa et al., 2016), pharmacological inhibition of αKG-dependent dioxygenases with DMOG impaired differentiation (Supplementary Fig. 5I), whereas prolonged supplementation with dimethyl-αKG enhanced *PAX6* expression (Supplementary Fig. 5J). Aspartate levels also increased transiently during the early progenitor stage before declining sharply by day 10 (Figure 4I), suggesting a stage-specific use of this metabolite.

Together with the early increased αKG levels and higher m+3/m+2 ratios in the progenitor stages, these findings suggest that early progenitors are not simply glycolytic but actively reroute glucose-derived carbon towards mitochondrial anaplerotic, biosynthetic and epigenetic pathways that support lineage commitment. As differentiation proceeds, pyruvate utilization progressively shifts towards PDH-mediated oxidation and mitochondrial energy metabolism.

### Pyruvate utilization determines the pace of neuronal maturation

Following the shift from PC- to PDH-mediated pyruvate utilization during differentiation, we next asked whether this transition is simply a consequence of differentiation or actively contributes to neuronal maturation. Because pyruvate partitioning between oxidative and anaplerotic pathways can be dynamically regulated to support distinct cellular functions (Elia et al., 2022), we pharmacologically manipulated mitochondrial pyruvate metabolism to test its role in neuronal differentiation.

During the early progenitor stages, when PC activity predominates, we activated the PDH complex with dichloroacetate (DCA), an inhibitor of pyruvate dehydrogenase kinases (PDKs), to prevent PDH phosphorylation and inactivation (Guo et al., 2023, Mathieu and Ruohola-Baker, 2017, Stacpoole et al., 2008, TeSlaa et al., 2016). Dose and timing optimization identified a 6-day treatment window from day 4 to day 10 as the most effective condition (Supplementary Fig. 6A–D). Although DCA treatment did not alter the m+3/m+2 labelling ratio (Figure 5B), it redirected glucose-derived carbon towards mitochondrial metabolism, as indicated by reduced lactate production and increased labelling of citrate and malate (Figure 5C). This suggests that the m+3/m+2 ratio does not fully capture changes in pyruvate utilization under these conditions, likely due to isotopic redistribution during subsequent TCA cycle turns. Importantly, PDH activation promoted neuronal maturation, resulting in increased expression of the neuronal marker NEFL and the motor neuron maturation markers ISL1 and ChAT(Figure 5D). Moreover, direct inhibition of PC with ZY-444 (Supplementary Fig. 6A–D), also modestly increased the expression of neuronal maturation markers (Supplementary Fig. 6E–F). Similar effects were observed with a different cell line (Supplementary Fig. 6G–H).

**Figure 5:**
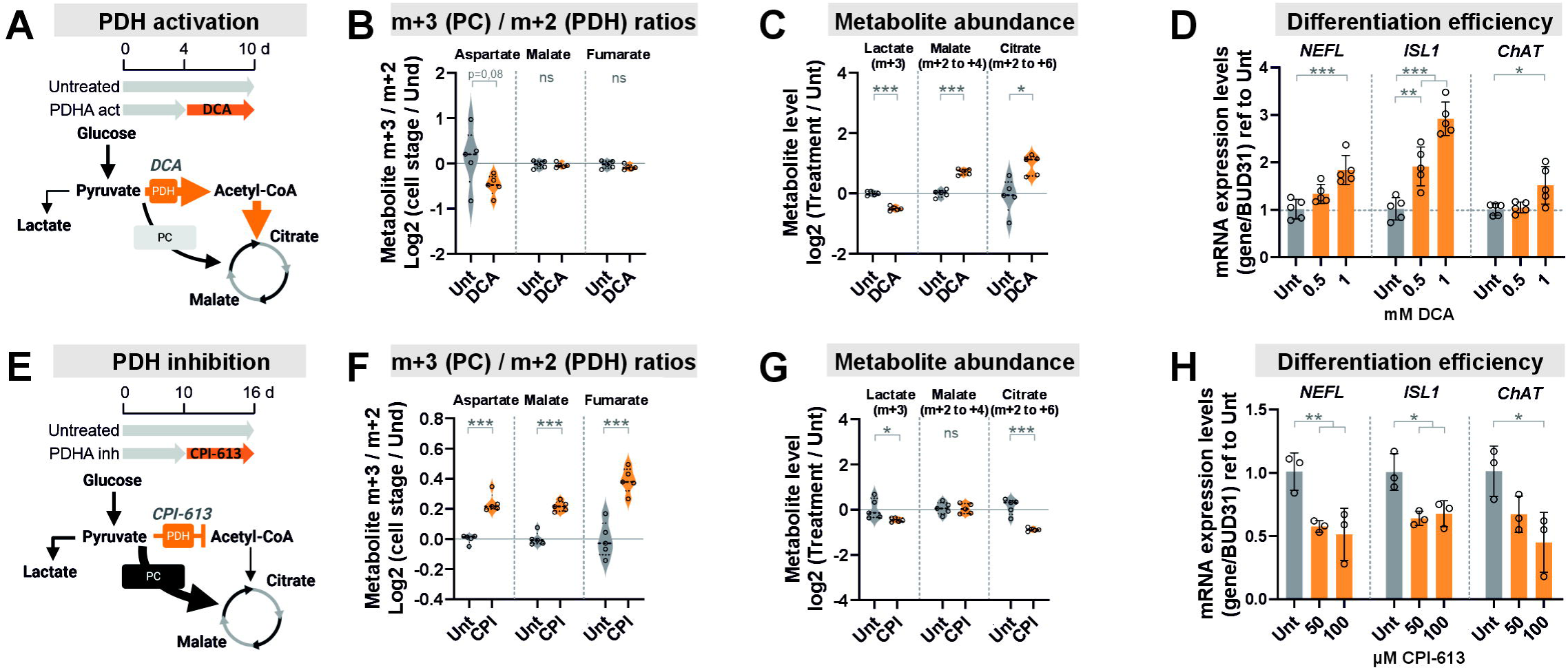
Mitochondrial pyruvate metabolism influences the pace of neuronal maturation. **A)** Cartoon schematic of the PDH activation strategy during progenitor stages. Created in BioRender. Torregrosa, R. (2026) https://BioRender.com/jqyagdk. Dichloroacetate (DCA, 0.5–1 mM) was added to the culture medium from day 4 to day 10, followed by metabolomics analysis. DCA activates PDH by inhibiting PDKs (Mathieu and Ruohola-Baker, 2017, Stacpoole et al., 2008, TeSlaa et al., 2016). **B)** Metabolite ratios of m+3 (reflecting pyruvate carboxylase–mediated anaplerosis) to m+2 (reflecting PDH-mediated entry) were used to infer pyruvate utilization following 1 mM DCA treatment. Lower ratios indicate a relative preference for PDH flux. **C)** Relative abundance of ¹³C isotopologues (as indicated below each metabolite name) in pyruvate-derived products following 1 mM DCA treatment. **D)** Differentiation efficiency following PDH activation, assessed by qPCR analysis of the expression levels of neuronal maturation markers (NEFL, ISL1, and ChAT). Values are presented as mean ± standard deviation and normalized to untreated (Unt) cells**. E)** Cartoon schematic of the PDH inhibition strategy during maturation stages. Created in BioRender. Torregrosa, R. (2026) https://BioRender.com/jqyagdk. CPI-613 (50–100 µM) was added to the culture medium from day 10 to day 16, followed by metabolomics analysis. CPI-613-mediated PDH inhibition reduces pyruvate-to-acetyl-CoA utilization, potentially increasing reliance on alternative anaplerotic pathways such as pyruvate carboxylase. **F)** Metabolite ratios of m+3 (reflecting pyruvate carboxylase–mediated anaplerosis) to m+2 (reflecting PDH-mediated entry) were used to infer pyruvate utilization following 100 µM CPI-613 treatment. Lower ratios indicate a relative preference for PDH flux. **G)** Relative abundance of ¹³C isotopologues (as indicated below each metabolite name) in pyruvate-derived products following 100 µM CPI-613 treatment. **H)** Differentiation efficiency following PDH inhibition, assessed by qPCR analysis of the expression levels of neuronal maturation markers (NEFL, ISL1, and ChAT). Data are presented as mean ± standard deviation and normalized to untreated (Unt) cells. Data are shown either as relative values (normalized to 1) or as log□ -transformed values, as indicated. In all cases, cell line: iPSC1; n = 3-6 technical replicates (wells). Statistical significance was defined as p < 0.05 (*), p < 0.01 (**), or p < 0.001 (***). Pairwise comparisons were performed using a t-test. Multiple group comparisons were conducted using one-way ANOVA followed by Dunnett’s multiple comparison test against the untreated control.

To determine whether the shift towards PDH-dependent utilization of pyruvate is required for maturation, we treated cells with CPI-613, an inhibitor of the PDH and α-ketoglutarate dehydrogenase (αKGDH) complexes. Dose and timing optimization identified a 6-day treatment window from day 10 to day 16 as the most effective condition (Supplementary Fig. 6I–L). CPI-613 treatment increased the m+3/m+2 labelling ratio (Figure 5E,F), indicating a relative shift towards PC-mediated pyruvate utilization. This was accompanied by reduced citrate labelling and lower lactate production (Figure 5G), consistent with altered pyruvate partitioning between oxidative and anaplerotic pathways. Importantly, this metabolic rewiring was sufficient to impair neuronal maturation, resulting in reduced expression of differentiation and maturation markers (Figure 5H).

These findings indicate that the developmental transition from PC- to PDH-dependent pyruvate utilization is not merely a consequence of neuronal differentiation but actively contributes to neuronal maturation. Notably, PDH inhibition increased the relative contribution of PC-mediated pyruvate metabolism, suggesting that PC-dependent anaplerosis can partially compensate when oxidative pyruvate metabolism is restricted in neurons.

## DISCUSSION

Mitochondrial remodelling is increasingly recognized as an integral part of cell fate transitions, yet the temporal organization of these organelles during human motor neuron differentiation remains poorly understood. Our results provide a temporally resolved reference resource integrating mitochondrial characterization, proteomics, metabolic profiling, and isotope tracing across defined stages of human iPSC-derived motor neuron differentiation, enabling the evaluation of mitochondrial maturation in stem cell-derived neuronal and disease models. These data reveal that in human motor neurons mitochondrial remodelling does not follow a straightforward progression from a glycolytic pluripotent state to a more OxPhos-dependent neuronal state. Instead, differentiating cells enter a prolonged intermediate state characterized by selective reorganization of mitochondrial content, quality-control pathways and metabolism before acquisition of a mature neuronal phenotype. By combining temporal resolution with multi-omics analysis, our study provides a framework for understanding how mitochondrial abundance, composition and function are coordinated during human motor neuron development.

Our findings uncover a distinct mitochondrial and metabolic state at the neural progenitor stage, revealing a broader principle that developmental progression can be structured by metabolic transitions that are invisible to traditional endpoint analyses. Mitochondrial abundance declined progressively during early differentiation and reached its lowest point around the progenitor stage before recovering during neuronal maturation. Similar transient reductions in mitochondrial content have been reported during differentiation toward neuroectodermal, mesodermal and endodermal lineages (Birket et al., 2011, Lees et al., 2018, Lv et al., 2022, Mostafavi et al., 2021). However, our temporal analysis demonstrates that this mitochondrial-low state persists far beyond pluripotency exit and represents a prolonged developmental phase during neurogenesis rather than a brief transitional event. Importantly, mitochondrial depletion was not accompanied by a uniform reduction of all mitochondrial functions. While mtDNA and the proteome in most mitochondrial pathways declined, certain mitochondrial proteins remained stable or followed independent trajectories, indicating that mitochondrial remodelling involves selective reorganization rather than global suppression of mitochondrial activity.

Although PSC stem cells rely predominantly on glycolysis and possess limited respiratory capacity compared with differentiated cells, mitochondria can contribute to biosynthetic, redox and signalling functions that are essential for maintenance of pluripotency (Arnold et al., 2022, Bukowiecki et al., 2014, Carey et al., 2015, Gu et al., 2016, Moussaieff et al., 2015). The subsequent reduction in mitochondrial abundance during differentiation therefore likely reflects a recalibration of the mitochondrial network capacity rather than a simple expansion from the starting state at pluripotency. Although selective mitochondrial turnover could contribute to the removal of dysfunctional organelles, mtDNA heteroplasmy levels remain remarkably stable throughout differentiation in our system (Kenvin et al., 2022), arguing against a strong selection based on mtDNA genotype. Instead, mitochondrial turnover may represent a developmental strategy to replace the mitochondrial network associated with the pluripotent state with one better suited to the metabolic requirements of neural progenitors. In sum, these findings support a biphasic model in which mitochondrial depletion precedes the expansion and maturation of the neuronal mitochondrial network.

Our data further suggest that mitochondrial depletion during motor neuron differentiation is an active and regulated process rather than a passive consequence of developmental progression. Mitochondrial fragmentation coincided with changes in autophagy- and lysosome-associated pathways, while inhibition of lysosomal acidification attenuated mitochondrial depletion. These features are consistent with programmed mitophagy described during developmental metabolic remodelling in different contexts (Boya et al., 2018, Naik et al., 2019, Rodger et al., 2018, Twig et al., 2008), suggesting that mitochondrial depletion is part of the developmental programme accompanying neural progenitor specification. Notably in motor neurons, this process occurred independently of NIX. Despite its established role in developmental mitochondrial clearance and neuronal remodelling (Esteban-Martinez et al., 2017, Ordureau et al., 2021, Sandoval et al., 2008, Schweers et al., 2007), genetic ablation of NIX, either alone or together with BNIP3, failed to alter mitochondrial abundance or differentiation in human motor neurons. These findings indicate that developmental mitochondrial turnover during motor neuron differentiation is mechanistically distinct from established NIX-dependent paradigms. The distinct temporal profiles of autophagy-associated proteins further suggest that this remodelling is not driven by a single coordinated autophagic programme and may instead reflect a restricted window of developmental macroautophagy.

The most striking metabolic feature of the progenitor state was a preferential routing of mitochondrial pyruvate. Despite reduced mitochondrial abundance, isotope tracing demonstrated substantial mitochondrial metabolic activity throughout early differentiation. This pattern identifies the progenitor stage as a specialized metabolic state rather than simply an immature version of the later mature neuron. PC-dependent anaplerosis replenishes TCA-cycle intermediates that can support biosynthetic demands associated with proliferation and lineage progression (Arnold and Finley, 2023, Inigo et al., 2021). Thus, preferential PC utilization may allow a reduced mitochondrial pool to sustain biosynthetic functions while maintaining substantial glycolytic activity. As cells matured into neurons, PDH activation increased together with the restoration of mitochondrial content and OxPhos subunits. This transition is consistent with the energetic demands of mature neurons and their dependence on mitochondrial pyruvate oxidation (Acin-Perez et al., 2023, Gray et al., 2014, Yang et al., 2021, Zheng et al., 2016).

Importantly, pharmacological manipulation of pyruvate utilization altered motor neuron differentiation efficiency, suggesting that the PC-to-PDH transition is not merely a marker of developmental progression but may contribute functionally to lineage maturation. Similar links between mitochondrial metabolism and neural cell state transitions have been reported during cortical neuron differentiation, cortical neuron maturation and adult neural stem cell activation (Danacikova et al., 2025, Iwata et al., 2023, Petrelli et al., 2023). Together with recent evidence that glucose metabolism can directly influence developmental signalling pathways (Stapornwongkul et al., 2025, Villaronga-Luque et al., 2025), these findings support the concept that metabolism actively contribute to developmental progression rather than simply responding to changing energetic demands. The mechanisms linking pyruvate partitioning to neuronal differentiation remain unclear but may involve altered availability of metabolites such as acetyl-CoA and α-Kg, both established regulators of chromatin state and cell fate (Moussaieff et al., 2015, TeSlaa et al., 2016).

Overall, our study identifies motor neuron differentiation as a process involving a transient mitochondrial remodelling state characterized by mitochondrial depletion, lysosome-associated turnover and PC-favoured metabolism before acquisition of a mature oxidative neuronal phenotype. By resolving the timing and coordination of these transitions, our findings provide a framework for understanding how mitochondrial dysfunction may disrupt human neurodevelopment and expose stage-specific vulnerabilities obscured at terminal endpoints.

## MATERIAL AND METHODS

### Experimental model

#### Ethical permissions

Human patient stem cell research was approved by the Coordinating Ethics Committee of the Helsinki and Uusimaa Hospital District (Nr 95/13/03/00/15). Animal experiments (teratoma assay) were approved by the Animal Welfare Committee of Southern Finland ESAVI-9734-2021. NOD-*scid*-gamma (NSG; 005557; The Jackson Laboratory) mice were housed under a 12 h light–dark cycle with food and water available ad libitum.

### Cell lines

This study included healthy human iPSC reprogrammed from human foreskin fibroblasts (HFF): iPSC1 (HEL24.3; UHi006-A, RRID:CVCL_9T96; Sendai virus; newborn donor), as well as iPSC derived from human skin fibroblasts: iPSC2 (HEL62.1; UHi021-B, RRID:CVCL_D0N7; Sendai virus; 57-year-old donor) and iPSC3 (HEL11.4; UHi007-B, RRID:CVCL_JE79; retrovirus; 83-year-old donor), all generated at the Biomedicum Stem Cell Center (Helsinki). The corresponding parental fibroblasts for each iPSC line were included in the study (Fibro 1 / iPSC 1, Fibro 2 / iPSC 2, Fibro 3 / iPSC 3). Human iPSC Kolf_2-C1 (WTSIi018-B-1) was also used. Moreover, this work also involved human embryonic stem cell (ESC) lines: ESC FES21 (UHe001-A, RRID:CVCL_B132), ESC FES29 (UHe003-A, RRID:CVCL_B134), both generated at the Biomedicum Stem Cell Center (Helsinki), and ESC H1 (WAe001-A, WiCell). All the iPSC clones had comparable morphology and expressed similarly the pluripotency markers. Experiments were done with cells from passage numbers between 20 and 45 (iPSC), 20 and 60 (hESC) and 2 and 10 (Fibro), depending on the cell line.

### Cell culture

Human PSC were cultured at 37□, in a humidified atmosphere, normoxia and 5% CO_2_; and maintained on Matrigel-coated plates (Corning, #CLS354234) with Essential 8 medium (Gibco, #A1517001) supplemented with E8-supplement, or Essential 8 Flex (Gibco, # A2858501) during weekends. iPSC were passed once or twice a week with 0.5 mM EDTA in PBS when 60% confluent. Fibroblasts were maintained on DMEM High-Glucose (Lonza #12-614F) supplemented with 2mM L-Glutamate (Life Technologies #250300081) and 10% FBS (Life Technologies #10270106).

To evaluate the effect of culture conditions on mitochondrial content, iPSC were cultured either on Mitomycin-C–inactivated mouse embryonic fibroblasts (MEFs) (Gibco, #A24903) or in conditioned medium collected from MEFs (MEF-CM). MEFs were seeded on 0.1% gelatin-coated plates and cultured as previously described for human skin fibroblasts. Two hours before adding the iPSC, the plates were washed once with PBS before adding ESC medium (KnockOut DMEM (Gibco, #10829018) supplemented with 20% KnockOut Serum Replacement (Gibco, # 10828010), 0.1 mM non-essential amino acids, 2 mM L-glutamine, 1 mM sodium pyruvate, 0.1 mM β-mercaptoethanol, and 8 ng/mL FGF2). iPSC was then thawed or passed directly onto the MEF layer. For feeder-free conditions, MEF-CM was prepared by collecting ESC medium without FGF2 after 24 hours of incubation on inactivated MEFs. Prior to feeding ESCs, MEF-CM was supplemented with 8 ng/mL FGF2. Medium was changed daily.

### iPSC-Motor Neuron differentiation

iPSC were differentiated into motor neurons using a dual-SMAD inhibition protocol (Guo et al., 2017, Maury et al., 2015). Neuronal basal medium was prepared as 1:1 of DMEM/F-12 (Gibco #31331-028) and Neurobasal® (Gibco #21103-049), and supplemented with N2 (Gibco #17502048), B-27 (Gibco # 17504044), 0.1mM L-ascorbic acid (Santa Cruz #sc-394304) and 100µg/mL primocin (InvivoGen, #ant-pm-05). To start neural induction, iPSC were dissociated using EDTA and plated on ultra-low attachment dishes (Nunclon Sphera 6-well Plate, 174932 ThermoFisher). On day 0, basal medium was supplemented with 3 μM Chir-99021 (Selleckchem # S1263), 40 μM SB431542 (Millipore # 616461), 0,2 μM LDN-193189 (Sigma # SML0559) and 5 μM Y-27632 (Selleckhem # S1049). The medium was replaced the next day. From day 2 to 6, media was changed to basal medium supplemented with 0.1 μM retinoic acid (ThermoFisher #044540.04) and 0.5 μM SAG (Millipore # 566660). On day 7, medium was replaced with neuronal basal medium supplemented with retinoic acid and SAG, 10ng/ml BDNF (Peprotech #450-02) and GDNF 10ng/ml (Peprotech # 450-10). On day 9, medium was replaced as before with 20 μM DAPT (Calbiochem #565770). On day 10, the motor neuron spheres were dissociated into single cells with Accumax (Invitrogen 00-4666-56), and motor neuron progenitors were plated on 50 μg/ml poly-D-lysine (Merck Millipore #A-003-E) and laminin 10 μg/ml (Sigma-Aldrich #L2020) coated plates at 70 000 cells / cm^2^. On day 11, fresh medium was added to the wells. On day 14, half of the medium was replaced with basal medium supplemented only with DAPT, BDNF and GDNF. From day 16 on, half of the culture medium was replaced with basal medium supplemented with 10 ng/ml BDNF, GDNF and CNTF (Peprotech #450-13). All the experiments were performed on day 30 mature motor neurons, unless otherwise indicated. Occasionally, “flat cells” appear in the motor neuron culture, however, we carefully monitored that the wells used in this study did not contain flat cells to avoid confounding results.

### iPSC-Cardiomyocytes differentiation

Stem cells were grown to near confluency, after which the dishes were washed once with DMEM/F-12 (Gibco #31331-028). One mL of RPMI (Gibco, #12004997) medium supplemented with B-27 minus insulin (Gibco, #15944204) and 6 μM Chir-99021 (Selleckchem # S1263) was then added to each well. The following day, an additional 1 mL of medium was added without removing the previous medium. On day 2, the medium was aspirated and replaced with 1 mL of the same basal medium supplemented with 5 μM IWR-1 (Sigma, #I0161). On day 3, an additional 1 mL of medium was added. On days 4 and 5, the medium was replaced with basal medium without supplements. Beginning on day 6, the basal medium was supplemented with B-27 with insulin (Gibco, #17504044). From day 9 onward, the same basal medium without glucose was supplemented with 5 mM L-lactate (Sigma, #L7022). Samples were collected on day 14.

### iPSC-pancreatic cells differentiation

Near-confluent plates of stem cells were dissociated using EDTA and seeded onto new Matrigel in Essential 8 medium supplemented with 5 μM ROCK inhibitor Y-27632 (Selleckchem; #S1049). Cells were plated at a density of 0.22 million cells/cm² to achieve confluence. Definitive endoderm induction and subsequent differentiation into pancreatic progenitors were performed following a well-established protocol up to stage 5, corresponding to 14 days of differentiation (Balboa et al., 2022)

### iPSC-embryoid bodies formation

iPSC were dissociated using EDTA and seeded into ultra-low-attachment dishes (Thermo Fisherm #174932). Cells were cultured in Essential 6 medium (Gibco, #A151640), allowing spontaneous differentiation and formation of embryoid bodies. Media was changed every other day and samples were collected on day 14.

### Teratoma assay

To perform teratoma assay, human induced pluripotent cells (iPSC1, iPSC2 and ESC H1) were cultured and ∼ 60% confluent dishes of human PSC were dissociated using EDTA and seeded into ultra-low-attachment plates (ThermoFisher #174932) in Essential 8 medium supplemented with 5 μM ROCK inhibitor Y-27632 (Selleckchem, #S1049) and 100 µg/mL Primocin (InvivoGen, #ant-pm-05). Stem cells formed spheres overnight on an orbital shaker. Approximately 1,000 aggregates were packed in PE-50 tubing. Under isoflurane anesthesia, the aggregates were grafted with Hamilton syringe (PN: 81341) under the kidney capsule of immunocompromised (NSG) mice. Buprenorphine (0,05-0,1 mg/kg) and carprofen (5 mg/kg) were used as analgesics for 3 consecutive days. Animals were sacrificed, and teratomas were collected at day 10, week 5, and week 6 (one animal died before the scheduled collection time).

### Genome editing

CRISPR/SpCas9 technology was used to generate NIX and BNIP3 knock-outs in iPSC 1 line. The crRNA sequences were designed by using online analysis tools Benchling (https://benchling.com/crispr) and CRISPOR (version 5.0; http://crispor.tefor.net/) and IDT CRISPR/ Cas9 design checker (https://www.idtdna.com/). Two different crRNAs were designed to introduce a deletion in the target genes. For NIX, a deletion in exon 2 was introduced in p.63-75, (c.187-223) (Wilhelm et al., 2022). For BNIP3, a deletion was introduced in exons 2-3 in p.40-88 (c.197-264) (Sulkshane et al., 2021). To perform the genome editing, each crRNA (IDT, CRISPR-CAS9 crRNA) and Atto™550 labelled tracRNA (IDT, #1075927) were resuspended with Nuclease-free Duplex buffer (IDT, #11-01-01-01) and incubated together to form 100µM of final gRNA duplex. Each gRNA was later combined with 10µg of recombinant SpCas9 nuclease (IDT, #1081058) to form the ribonucleoprotein (RNP) complex. A total of 3µl of RNP were delivered by electroporation using the Neon Transfection System (ThermoFisher, USA) to 2 million Hel 24.3 cells, dissociated with Accutase (ThermoFisher, #A11105-01), together with 2 μl of 100 μM electroporation enhancer (IDT, #1075915). Limited dilution cloning followed the electroporation step. Briefly, 500-1000 TrypLE Select (Thermo Scientific, # 12563011) dissociated cells were plated into 10 cm tissue-culture dishes coated with Matrigel and grown E8-flex medium supplemented with 1X CloneR (STEMCELL Technologies, #05888). Following 48 hours incubation, cell culture medium was replaced with 10 mL of E8-Flex medium supplemented with E8-Flex medium supplement and changed every 48 hours for 8 days until colony picking. Knockout colonies were validated using agarose gel electrophoresis, followed by western blot to confirm the absence of the protein of interest. For NIX knockout generation, the guides used were the following, sense: TTGTGGATGGAGGATGAGGA, and anti-sense: AGACATGGAGAAGATTCTTT. For BNIP3 knockout generation the guides used were, sense: GGAGAGAAAAACAGCTCAC and anti-sense: TCCATTATAAATAGAAACCG. For knockout verification, the primers used for NIX were the following: forward CCAACAGGTTCCTGGGTGGAGC, and reverse GTGGGCACTGAAGCACCTGTCA. And for BNIP3, forward: CAGCAGCAATGGCAATGATAA, and reverse: GTTACAGGGCTCATTCACTCTAG.

### Method details

#### Pharmacological treatments in cell culture

Lysosomal acidification was inhibited with 0.5 nM bafilomycin A1 (BafA1) (MedChem Express, #HY-100558) for 2 or 4 days starting from day 0. Supplementation with α-ketoglutarate was performed using 6 µM dimethyl-2-oxoglutarate (dm-αKG; Sigma, #349631) for 3, 6 or 10 days starting from day 0, whereas inhibition of αKG-dependent dioxygenases with 150-300 µM DMOG for 4 days starting from day 0. Activation of the PDH complex was achieved using 0.5-1 mM dichloroacetate (DCA) (Merck, #347795) and inhibition of pyruvate carboxylase with 0.625-1.25 µM ZY□ 444 (MedChem Express, HY-142870) for 6 days from day 4 to day 10. Inhibition of PDH was carried out using 50-100 µM CPI□ 613 (MedChem Express, HY-15453) for 6 days, from day 10 to day 16.

#### Total DNA extraction and mtDNA copy number analysis

Total cellular DNA (mitochondrial and nuclear) was isolated using proteinase K (ThermoFisher # E00491) and SDS lysis (10mM Tris pH 7.4, 10mM EDTA, 150mM NaCl and 0.4% SDS) followed by phenol:chloroform extraction and ethanol precipitation (Torregrosa-Munumer et al., 2019) using Ultrapure Phenol:Chloroform:Isoamyl Alcohol (ThermoFisher #15593031). To increase PCR efficiency and minimize variability arising from different DNA topologies, total DNA samples were gently sonicated using an ultrasonic processor (VibraCell, TAMRO) operated at a 50% duty cycle. Total DNA was first assessed by absorbance (QFX Fluorometer, DeNovix) to evaluate sample quality and subsequently quantified using a fluorescent die with the Qubit™ dsDNA Quantification Assay Kit (Invitrogen, #Q32850). Levels of mtDNA were quantified by qPCR targeting the mitochondrial MT-ND5 or MT-tRNA^LEU^ gene and normalized to the nuclear single-copy gene B2M. Reactions were performed on a CFX Real-Time System C1000 Touch (BioRad) with DyNAmo Flash SYBR Green qPCR Kit (ThermoFisher # F415L), using 0.5µM per primer and 2ng of DNA for nuclear DNA or 0.2ng of DNA for mtDNA reactions(Wong and Cortopassi, 2002). The following thermoprofile was used: 7 minutes at 95□ followed by 40 cycles of a 2-step amplification: denaturation at 95□ for 10 seconds and annealing and elongation at 60□ for 30 seconds. Each sample was measured by duplicate. Relative mtDNA copy number was calculated using the ΔΔCt method. The primers used were *B2M* (Forward: TGCTGTCTCCATGTTTGATGTATCT, Reverse: TCTCTGCTCCCCACCTCTAAGT), *MT-ND5* (Forward: AGGCGCTATCACCACTCTGTTCG, Reverse: AACCTGTGAGGAAAGGTATTCCTG) and MT-tRNA^LEU^ (Forward: CACCCAAGAACAGGGTTTGT, Reverse: TGGCCATGGGTATGTTGTTA).

To assess the variation between total DNA and mtDNA levels during the first 10 days of MN differentiation, total DNA was extracted as described above and divided into two fractions. One fraction was used to quantify total DNA by fluorescence using the Qubit™ dsDNA Quantification Assay Kit (Invitrogen, # C35011). The second fraction was diluted 1:50 for all samples and subjected to qPCR analysis using the mtDNA-specific primers. Comparable amounts of DNA were not standardized across assays; instead, each assay was performed using the raw extracted material. In this context, increases in total DNA reflect increases in cell number and proliferation, against which mtDNA levels may rise proportionally (or diverge) depending on mitochondrial dynamics.

#### RNA extraction and gene expression analysis

RNA was extracted using the NucleoSpin RNA Extraction Kit (Macherey-Nagel, #740955) and quantified with a spectrophotometer (QFX, DeNovix). A total of 40 ng RNA was reverse-transcribed using the Maxima First Strand cDNA Synthesis Kit (Thermo Fisher, #K1641), and 5 ng of the resulting cDNA were analyzed by qRT-PCR on a CFX Real-Time System C1000T (Bio-Rad) using SYBR Green Flash (ThermoFisher, #F415L). Each sample was measured by duplicate. Relative mRNA expression was calculated using the ΔΔCt method and normalized to β*-Actin* or *BUD31 (*BUD31 was selected based on proteomics data, as it showed minimal variation during the mid-stages of differentiation). Primer sequences are: *ACTA2* (Forward: CTATGCCTCTGGACGCACAACT, Reverse: CAGATCCAGACGCATGATGGCA), *ChAT* (Forward: GGAGGCGTGGAGCTCAGCGACACC, Reverse: CGGGGAGCTCGCTGACGGAGTCTG), *Hb9* (Forward: GTCCACCGCGGGCATGATCC, Reverse: TCTTCACCTGGGTCTCGGTGAGC), *ISL1* (Forward: GTTACCAGCCACCTTGGAAA, Reverse: GGACTGGCTACCATGCTGTT), *Nanog* (Forward: CTCAGCCTCCAGCAGATGC; Reverse: TAGATTTCATTCTCTGGTTCTGG), NEFL (Forward: CAAGACCCTGGAAATCGAAG, Reverse: TGAAACTGAGTCGGGTCTCC), *PAX6* (Forward: ATGGGCGGAGTTATGATACCTAC; Reverse: GGAACTTGAACTGGAACTGACA), *SOX17* (Forward: CGCCGAGTTGAGCAAGAT, Reverse: GGTGGTCCTGCATGTGCT), *Olig2* (Forward: ATGCACGACCTCAACATCGCCA, Reverse: ACCAGTCGCTTCATCTCCTCCA). As housekeeping genes, β*-actin* (forward: CCTGGCACCCAGCACAAT, reverse: GGGCCGGACTCGTCATAC) or *BUD31* (Forward: CATTCAGACACGGGACACCA, Reverse: ATGATGCGGCCCACTTCC) were used.

### Immunocytochemistry

Fibroblasts, iPSC, ESCs and iPSC-MN were cultured on cover glasses and fixed with 4% PFA for 15 min at room temperature and then permeabilized with PBS containing 0.2% Triton X-100 (FisherScientific #BP151-100) for 10 min. Cells were blocked with 5% protease-free BSA (Biowest # P6154) in 0.1% Tween20 (Sigma, #P1379) in PBS-T for 2 h in RT, and then were incubated overnight at +4°C in blocking buffer containing 5% bovine serum and primary antibodies: Hb9 (DSHB #81.5C10-s, 1:50), ISL1 (DSHB #39.4D5-s, 1:50), MAP2 (Abcam #5392, 1:1000), TUJ1 (Biolegend #801201, 1:1000) or SDHA (Abcam #ab14715, 1:2000). The following day, cells were washed with PBS-T and then incubated with the secondary antibody diluted 1:1000 for 1h at RT: AlexaFluor^TM^ 594 anti-rabbit IgG (A11012, Thermo Fisher), AlexaFluor^TM^ 488 anti-mouse IgG (A11008, Thermo Fisher), AlexaFluor^TM^ 488 anti-goat IgG (A11055, Thermo Fisher), DyLight 488 anti-chicken IgG (SA5-10070, Thermo Fisher). Cover glasses were applied on microscope slides with Vectashield DAPI (VectorLabs #H-1200) and imaged with Axio Observer Z1 (Zeiss) inverted fluorescence microscope. Quantification of Hb9- and ISL1/2-positive cells was performed manually using a total of 24–29 images, encompassing 2240 –3186 cells for Hb9 and ISL1/2, respectively.

### Transmission electron microscopy

Cells growing as monolayers were cultured on cover slips and fixed in 0.1 M NaCac buffer containing 2% glutaraldehyde for 30 minutes at room temperature. Fixed cells were processed by standard methods and imaged with Jeol JEM-1400 transmission electron microscopy (TEM) (Jeol Ltd., Tokyo, Japan) operating at 80 kV. Mitochondrial parameters were analyzed using Microscopy Image Browser (Belevich et al., 2016) from at least 20 independent, non-overlapping images per time point. Individual mitochondria were defined as separate objects, and measurements were averaged within each image. Image-level means were used as individual data points for statistical analysis. The total number of mitochondria analysed is indicated in the corresponding figure legend.

To compare day 0 (undifferentiated iPSC cultured as a monolayer) with day 2 neural progenitors (normally cultured in suspension as neurospheres), day 2 NMPs were dissociated with AccuMax for 5 minutes and seeded onto Matrigel-coated dishes. After allowing the cells to attach and recover for 5 hours in day 2 MN differentiation media, fixation was performed following the same protocol used for monolayer cultures. Mitochondrial DNA content and early differentiation markers (SOX2 and PAX6) did not differ between 2D and 3D conditions.

To compare neurospheres from day 2 to day 10, neural progenitor cells were culture in suspension as neurospheres following the standard differentiation protocol and then were fixed with 2% glutaraldehyde and 2% PFA in 0.1 M NaCac buffer pH 7.4, supplemented with 2 mM CaCl2, for 2 h, at RT. 1:10 volume of pellet:fixative was used. After this, fixative was changed to 2% PFA in 0.1 M NaCac buffer pH 7.4, supplemented with 2 mM CaCl2 and stored o/n at 4°C. Fixed spheroids were then Washed 2 x 5 min with 0.1 M NaCac pH 7.4 then osmicated with 1% OsO4 in 0.1 M NaCac for 1 hour, on ic. After this they were Washed 2 x 5 min with 0.1 M NaCac, pH 7.4 followed by a dehydration with 50%, 70%, 96%, 100% EtOH for 10 min and with acetone 3 x 10 min. Dehydrated spheroids were then Embedded into Epon (hard, TAAB) with 1:3 Epon:acetone 3 h, 3:1 Epon:acetone o/n, 100% Epon 3 h, 100% Epon 3 h and baked at +60°C, > 14 h. Spheroids were imaged and analyzed as described above.

### Western blot

Total protein was obtained by lysing cells in RIPA buffer (ThermoFisher # 89900) supplemented with 1x Halt™ protease inhibitor (Thermo Scientific #87786). Protein concentration was quantified using the BCA Protein Assay Kit (Pierce, #23225) according to the manufacturer’s instructions. Then, 10-15 µg of protein per sample were prepared in 4x Laemmli buffer (BioRad # 1610747), boiled for 5 minutes at 95°C and separated on 4–20% Mini-PROTEAN® TGX™ Precast Protein Gels (BioRad #4561094). Proteins were transferred to a Trans-Blot Turbo Mini 0.2 µm Nitrocellulose (BioRad #1704158) using the Trans-Blot Turbo transfer system (Bio-Rad). Membranes were blocked with 5% milk (or 3% BSA for phosphorylated proteins) in TBS-T for 1 hour at room temperature, followed by overnight incubation with primary antibodies prepared in 3% BSA in TBS-T: ATP5A (Abcam #ab14748, 1:1000), BNIP3 (Abcam, # ab109362, 1:1000), COXII (Abcam, #ab79393, 1:1000), DNAJA3 (TID1), DRP1 (CellSignaling, #5391, 1:1000), DRP1 s616 (Cell Signaling, #3455, 1:1000), DRP1 s637 (Abcam, #ab193216, 1:1000), GAPDH (Cell Signaling, # 2118, 1:1000), Histone 3 (Cell Signaling, # 4499, 1:1000), ((Abcam, #ab181024, 1:1000), HSP60 (Santa Cruz, #sc-1052, 1:200), MT-CO1 (Abcam, #ab14705, 1:1000), Nanog (Cell Signaling, #4903S, 1:2000), NDUFA9 (Abcam, #ab14713, 1:2000), NIX (Cell Signaling, #12396S, 1:1000), COX7A2L (Proteintech #11416-1-AP, 1:500), LC3B (Abcam, #ab51520, 1:1000), LDHA (Cell Signaling, #3582, 1:1000), SDHA (Abcam #ab14715, 1:2000), SDHB (Abcam #ab14714 1:1000), SOX1 (R and D Syst, #MAB3369, 1:1000), SQTM1 (p62) (Abnova, #H00008878-M01, 1:5000), PDHA1 (Abcam #ab110330, 1:2000), PDHA1 S293 (Abcam #ab177461, 1:2000), TOM20 (Santa Cruz, #sc1-17764, 1:500), TOM40 (Santa Cruz, #sc-11414, 1:1000), UQCRC2 (Abcam, #ab14745, 1:1000; Proteintech #14742-1-AP, 1:1000), VDAC1/3 (porin) (Abcam, #ab14734, 1:2000), Vinculin (Sigma, #V9264-25UL, 1:10000). The next day, after three washes of 20 minutes each with TBS-T, membranes were incubated with secondary antibodies for 1 hour: Peroxidase Goat Anti-Mouse IgG (Jackson #115-035-146, 1:5000), Peroxidase Goat Anti-rabbit IgG (Jackson #111-035-003, 1:5000), all prepared in 3% BSA in TBS-T. Following another three 20-minute washes with TBS-T, the luminescent signal was detected using ECL reagent (Thermo Scientific), and membranes were imaged with the ChemiDoc XRS+ (Bio-Rad). Band intensities were quantified using Image Lab Software (Bio-Rad). For the analysis, total protein was quantified by Ponceau staining using the Ponceau S Solution (Fisher Scientific, #33427.01). Alternatively, when applicable, the cytosolic signal of vinculin was used as a loading control for total protein normalization, and VDAC1/3 was used for mitochondrial protein normalization. For phosphorylation studies, cell pellets were processed immediately after collection to preserve phosphorylation status.

### Blue-Native PAGE

Cells were dissociated using AccuMax (A7089, Sigma-Aldrich), counted, and assessed for viability. For mitochondrial enrichment, 5 million iPSC, 10 million cells from days 4 and 10, and 2 million cells from day 30 were collected by centrifugation and resuspended in 200 µL of PBS. An equal volume (200 µL) of 4 mg/mL digitonin was added and samples were incubated on ice for 10 min to solubilize mitochondrial membranes. Subsequently, 1 mL of ice-cold PBS was added, and the samples were centrifuged at 10,000 × *g* for 5 min at 4°C. The supernatant was discarded, and the resulting mitochondria-enriched pellet was washed once with 1 mL of ice-cold PBS. After a second centrifugation at 10,000 × *g* for 5 min at 4°C, the supernatant was removed and the pellet was stored at −80°C until further analysis. Blue Native PAGE (BN-PAGE) was performed as previously described (Fernandez-Vizarra and Zeviani, 2021).

### Oxygen consumption and ATP production rates

Mitochondrial oxygen consumption (OCR) and extracellular acidification rate (ECAR) in iPSC and iPSC-MN was determined using Seahorse XF96 Extracellular Flux Analyzer and Mitostress test (Agilent). Fibroblasts were seeded at 20.000 cells per well and PSC at 25.000 cells per well on Matrigel-coated XF96 Seahorse plates (Matrigel; Corning) in the presence of 10 μM ROCK inhibitor Y□ 27632 (Selleckchem, #S1049) the day before the experiment. (Zhang et al., 2012). The following day, the medium was replaced one hour before measurement with Seahorse XF Base Medium (Seahorse XF Base Medium; Agilent #103335-100) supplemented with 1 mM pyruvate, 2 mM glutamine, 17.5 mM glucose, and 5 mM HEPES, adjusted to pH 7.4. OCR was recorded three times under baseline conditions and three times following sequential injections of 2 µM oligomycin (Sigma, #O4876), 2 µM FCCP (Sigma, #C2920), and 1 µM rotenone (Sigma, #R8875) plus antimycin-A (Sigma, #A8674). For iPSC-MNs, 20,000 cells were plated on poly-D-lysine/laminin-coated XF96 Seahorse plates on day 10 of differentiation and maintained until day 15 or day 30 following the MN differentiation protocol. At day 15 or 30, OCR measurements were performed as for iPSC, using 2 µM oligomycin, 5 µM FCCP, and 1 µM rotenone plus antimycin-A. Data were normalized to DNA content (as a proxy for cell number) using the CyQUANT Cell Proliferation Assay Kit (Invitrogen, # C35011). Only wells with a homogeneous monolayer of cells were included in the analysis.

### High-throughput methods

#### Metabolic profiling

For metabolic profiling experiments, the iPSC 1 cell line was used. For the metabolic tracing, undifferentiated (day 0) and iPSC-MN (days 4, 10, 16 and 30) were incubated with 21.25□ mM U-^13^C glucose (Cambridge Isotope Laboratories #CLM-1396) and 2 mM glutamine in DMEM-F12 (w/o glucose, Biowest, #L0091) for 30 or 180 minutes. Concentration of glucose and glutamine were comparable to routine neural culture conditions. For metabolites extraction, cells were washed once with cold PBS and 150 µl / million cells of ice-cold 20:80 H_2_O/acetonitrile (Sigma # 34851) was added. Cells were scraped, collected, and vortexed for 5 seconds. Finally, extracted samples were centrifuged at 13,000g for 10 minutes at 4°C, and the supernatant was stored at -80°C until analysis. For each time point, 6 separate wells were used as technical replicates. Steady state metabolomics experiment was done similarly, but with unlabeled substrates and only included 3 replicates (wells).

Extracted metabolites were analyzed with a Thermo Q Exactive Focus Quadrupole Orbitrap mass spectrometer coupled with a Thermo Dionex UltiMate 3000 HPLC system (ThermoFisher Scientific). The HPLC was equipped with a hydrophilic ZIC-pHILIC column (150□×□ 2.1□mm, 5□μm) with a ZIC-pHILIC guard column (20□×□ 2.1□mm, 5□μm, Merck Sequant). A 5□µl sample was injected into the LC–MS instrument after quality controls in randomized order having every tenth samples as blank. A linear solvent gradient was applied in decreasing organic solvent (80–35%, 16□min) at 0.15□ml□min–1 flow rate and 45□°C column oven temperature. Mobile phases were aqueous 200□ mmol per litre ammonium bicarbonate solution (pH□ 9.3, adjusted with 25% ammonium hydroxide), 100% acetonitrile and 100% water. Ammonium bicarbonate solution was kept at 10% throughout the run, resulting in a steady 20□mmol per litre concentration. Metabolites were analyzed using a mass spectrometer with a heated electrospray ionization source using polarity switching and the following settings: resolution of 70,000 at m/z of 200; spray voltages of 3,400□V for positive and 3,000□V for negative mode; sheath gas of 28 arbitrary units (AU) and auxiliary gas of 8□AU; vaporizer temperature of 280□°C; and ion transfer tube temperature of 300□°C. The instrument was controlled using Xcalibur 4.1.31.9 software (Thermo Scientific). Metabolite peaks were confirmed using commercial standards (Sigma-Aldrich). Data quality was monitored throughout the run using an in-house quality control cell line extracted in a similar manner to other samples.

Metabolite peaks areas were exported from TraceFinder and analyzed in R and Python. Peak areas were corrected by subtracting the areas observed in blank samples (extraction buffer) and normalized to the total metabolite intensity (TMI) of each sample. For differential abundance analysis, TMI-normalized metabolite abundances were subsequently quantile normalized to account for substantial variation in TMI and protein concentration across differentiation stages. Missing values were replaced with zeros while recording the number of missing observations for each metabolite. Differential abundance between genotypes was assessed using limma (Ritchie et al., 2015), fitting linear models with lmFit and applying empirical Bayes moderation with eBayes. Metabolites with FDR-corrected p-values < 0.01 were considered differentially abundant if they contained less than 3 missing values. Labeled fractions were obtained after combining the peak areas of the labeled (m+x to m+y) isotopologues unless otherwise indicated.

### Proteomics

For total lysate proteomics, the iPSC 1 cell line was used. On days 0, 2, 4, 10, 16, 21 and 30, cells were washed once with PBS, and then lysed in 8 M urea in 50 mM Tris-HCl, pH 8.0 and stored at -80°C until analysis. Five separate wells were used as technical replicates.

Samples were reduced with 10 mM D,L-dithiothreitol and alkylated with 40 mM iodoacetamide and then digested overnight with sequencing grade modified trypsin (Promega). After digestion peptide, samples were desalted with a Sep-Pak tC18 96-well plate (Waters) and evaporated to dryness. Samples were dissolved in 0.1% formic acid and peptide concentration was determined with a NanoDrop device. The LC-ESI-MS/MS analysis was performed on a nanoflow HPLC system (Easy-nLC1200, ThermoFisher Scientific) coupled to the Orbitrap Fusion Lumos mass spectrometer (ThermoFisher Scientific) equipped with a nano-electrospray ionization source and FAIMS interface. Compensation voltages of -50 V and -70 V were used. Peptides were first loaded on a trapping column and subsequently separated inline on a 15 cm C18 column (75 μm x 15 cm, ReproSil-Pur 3 μm 120 Å C18-AQ, Dr. Maisch HPLC GmbH, Ammerbuch-Entringen). The mobile phase consisted of water with 0.1% formic acid (solvent A) or acetonitrile/water (80:20 (v/v)) with 0.1% formic acid (solvent B). A 120 min gradient was used to elute peptides (60 min from 5 % to 21 % solvent B followed by 48 min from 21 % to 36 % solvent B and in 6 min from 36% to 100% of solvent B, followed by 6 min wash stage with solvent B). Samples were analyzed by a data independent acquisition (DIA) LC-MS/MS method. Wash runs were submitted between each sample to reduce potential carry-over of peptides. MS data was acquired automatically by using Thermo Xcalibur 4.6 software (ThermoFisher Scientific). In a DIA method a duty cycle contained one full scan (400-1000 m/z) and 30 DIA MS/MS scans covering the mass range 400 - 1000 with variable width isolation windows.

Protein identifications were done using DirectDIA and label free quantifications were done with MaxLFQ using Spectronaut software (Biognosys; version 18.0.2) against Swiss-Prot (2023_01 Homo Sapiens) and Universal Protein Contaminant database (Frankenfield et al., 2022). Precursor and protein detection FDR cutoff was 0.01. Quantification was carried out at MS2 level with area under the curve within integration boundaries for each targeted ion. The protein peak areas were exported from Spectronaut and analysed in R. Missing values were replaced with zeros, while flagging the number of missing proteins. Protein peak areas were log1p-transformed and quantile normalized using voom (Ritchie et al., 2015). The differential expression between genotypes was analysed in R with limma (Ritchie et al., 2015) using lmFit with eBayes correction on the contrasts between genotypes. Proteins with FDR-corrected p-values < 0.01 were considered differentially expressed if they contained less than 3 missing values. Differentially expressed proteins were analysed for enrichment against Reactome, KEGG, DO, WP and GO databases using clusterProfiler (Yu et al., 2012). Figures were created with R ggplot2 and Python Matplotlib. Mitochondrial proteins were identified using MitoCarta 3.0 (Rath et al., 2021), while other organelle assignments were based on previously published compartment annotations (Ordureau et al., 2021).

The normalized and log-transformed expression values were rescaled per protein, and the expression levels were centered around the mean expressions. The centered and rescaled expressions were clustered using hierarchical clustering with Wald D2 method, from the computed Euclidian distances. The dendrograms were cut into 8 clusters for the analysis looking at all proteins.

Trajectory analyses were performed using non-imputed proteomics data. Proteins with at least 14 identified peptides across all samples, quantified in all five day-0 wells, and with a mean day-0 abundance greater than zero were retained. Protein abundances were log□-transformed and normalized to the mean abundance of each protein across the five day-0 wells. For each sample, normalized abundances were averaged across all proteins within a given subset, yielding five independent observations per time point. Temporal trajectories were analyzed using ordinary least-squares regression with HC3 heteroscedasticity-robust standard errors, including time point, protein subset, and their interaction. Global differences among trajectories were assessed using joint Wald χ² tests, followed by pairwise comparisons between each subset and a designated reference subset. P values were adjusted for multiple testing using the Benjamini–Hochberg false discovery rate method.

For heatmaps, proteins included in pathway-specific heatmaps were selected based on Reactome and KEGG pathway assignments, MitoCarta 3.0 annotations, or manual literature curation. Only proteins supported by at least 14 unique peptides across the dataset and detected in all five replicates of at least one differentiation stage were included. This filter was used to ensure reliable protein identification while retaining proteins that may be expressed only during specific stages of differentiation and therefore may be absent at other time points. Within each pathway, proteins were ordered according to their mean abundance at either day 10 or day 30, depending on the analysis. Missing values were replaced with a small constant (1 × 10□□) prior to protein-wise Z-score normalization.

### Statistics and Reproducibility

Data are presented as mean ± standard deviation in all graphs. P-values < 0.05 were considered statistically significant. Statistical analyses were performed using GraphPad Prism v10.1.2 (GraphPad Software). Differences between two groups were assessed using unpaired two-tailed t-tests (Welch’s correction when appropriate). For comparisons involving more than two groups, one-way ANOVA followed by Tukey’s or Dunnett’s multiple-comparison test was used, as indicated. For the initial characterization of mitochondrial parameters, three independent cell lines (biological replicates) were analyzed, each including at least three wells (technical replicates). For high-throughput experiments and downstream studies Follow-up experiments, the iPSC1 cell line was preferentially used including at least three technical replicates (wells), unless otherwise indicated.

## DATA AVAILABILITY

Steady state metabolomics and isotope tracing raw data has been deposited to Metabolomics Workbench (Sud et al., 2016) (steady state: ST005242; extracellular isotope tracing: ST005243; intracellular isotope tracing: ST004580; DCA and ZY-444 treatments: ST005241; and CPI-613: ST005240), and the normalized and blanked data can be found in Supplementary Table 1. Proteomics raw data has been deposited in MassiveDB (MSV000100301) and can be found together with the analyzed data as Supplementary Table 2. Source data for all figures can be found as Supplementary Table 3. R and Python code to process and analyze the high throughput data using standard pipelines is available upon request.

## CODE AVAILABILITY

R (version 4.5.1) and Python (version 3.11) code to process, analyze and visualize the high throughput data using standard pipelines is available upon request.

## ACKNOWLEDGEMENTS

We acknowledge the expert support by Miia Nissilä and Juhana Juutila. We thank Ras Trokovic from the Biomedicum Stem Cell Center (BSCC), University of Helsinki, for support with the stem cell work. We acknowledge Elena Herbers support for technical assistance with the Seahorse experiments. We also thank Steffi Goffart, Jaakko Pohjosmäki, Anu Hangas and Koit Aasumets (University of Eastern Finland, Finland), and Fumi Suomi (University of Helsinki) for their efforts in testing and optimizing additional experimental approaches. We thank the Electron Microscopy Unit of the Institute of Biotechnology, University of Helsinki, for providing laboratory facilities. Mass spectrometry analyses were performed at the Turku Proteomics Facility, supported by Biocenter Finland. This study has been supported by the Research Council of Finland Centre of Excellence in Stem Cell Metabolism (grants #312436, #336194, #353000) to H.T., P.K., V.H. and T.O., Sigrid Juselius Fundation to H.T., and Research Council of Finland (grants #340207 and #330641) to R.T-M. S. P-C. is a recipient of a Ramón Areces Postdoctoral Fellowship.

## AUTHORS CONTRIBUTIONS

Conceptualization, R.T-M. and H.T.; Methodology, R.T-M., J.T., D. S., J.K., J.S-V., S. P-C., R.E., J.P., E. F-V.; Investigation, all authors.; Writing – Original Draft, R.T-M., and H.T.; Writing – Review and Editing, all authors; Funding Acquisition, R.T-M., H.T., P.K., V.H., and T.O.; Supervision, R.T-M. and H.T.

## DECLARATION OF GENERATIVE AI AND AI-ASSISTED TECHNOLOGIES IN THE MANUSCRIPT PREPARATION PROCESS

During the preparation of this work, the authors used CurreChat (a secure, GDPR-compliant generative AI service developed at the University of Helsinki) to improve language and readability, and to assist with literature exploration and synthesis. All AI-generated content was reviewed, verified, and edited by the authors, who take full responsibility for the content and conclusions of the published article.

## MATERIALS AND CORRESPONDENCE

Requests for further information and resources should be directed to and will be fulfilled by Ph.D. Rubén Torregrosa-Muñumer

## SUPPLEMENTAL INFORMATION

Document S1. Supplementary Figures.

Table S2: Analyzed metabolomics data.

Table S3: Analyzed proteomics data.

Table S4: Source data for all figures.

