## Supplementary_Data for "Metabolic commitment to human motor neurons is characterized by mitochondrial depletion and pyruvate rewiring"

**Content:**

- **Supplementary Figure 1: coordinated remodeling of mtDNA content and the proteome during human motor neuron differentiation**

- **Supplementary Figure 2: pluripotent stem cells have a large number of mitochondria but it is immature.**

- **Supplementary Figure 3: mitochondrial depletion during the exit of pluripotency is driven by fragmentation and lysosomal acidification.**

- **Supplementary Figure 4: coordinated remodeling of mitochondrial OxPhos complexes during neuronal maturation.**

- **Supplementary Figure 5: metabolic shift from glycolysis-dominant early progenitors to enhanced OxPhos during neuronal maturation.**

- **Supplementary Figure 6: mitochondrial pyruvate metabolism influences the pace of neuronal maturation.**

- **Supplementary Figure 7: uncropped Western blotting images for main figures**

- **Supplementary Figure 8: uncropped Western blotting images for supplementary data figures**

- **Supplementary bibliography**

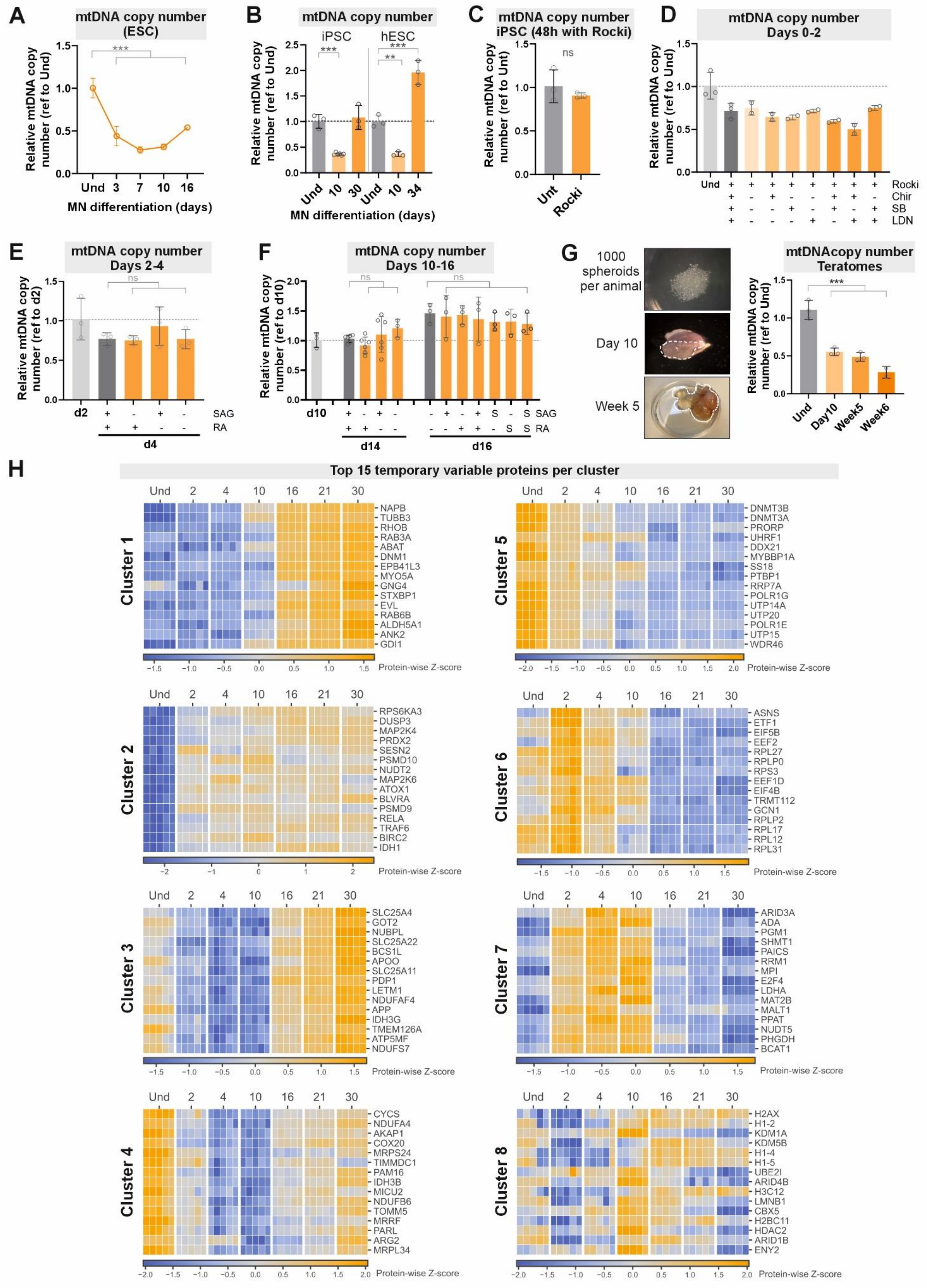
A
mtDNA copy number (ESC)
Relative mtDNA copy number (ref to Und)
MN differentiation (days)
Und 3 7 10 16
B
mtDNA copy number
iPSC
hESC
Relative mtDNA copy number (ref to Und)
Und 10 30 Und 10 34
MN differentiation (days)
C
mtDNA copy number iPSC (48h with Rocki)
ns
Relative mtDNA copy number (ref to Unt)
Unt Rocki
D
mtDNA copy number Days 0-2
Relative mtDNA copy number (ref to Und)
Rocki
Chir
SB
LDN
E
mtDNA copy number Days 2-4
Relative mtDNA copy number (ref to d2)
SAG
RA
d2
d4
F
mtDNA copy number Days 10-16
Relative mtDNA copy number (ref to d10)
ns
SAG
RA
d10
d14
d16
G
1000 spheroids per animal
Day 10
Week 5
mtDNAcopy number Teratomes
Relative mtDNA copy number (ref to Und)
Und Day10 Week5 Week6
H
Top 15 temporary variable proteins per cluster
Und 2 4 10 16 21 30
Cluster 1
NAPB
TUBB3
RHOB
RAB3A
ABAT
DNM1
EPB41L3
MYO5A
GNG4
STXBP1
EVL
RAB6B
ALDH5A1
ANK2
GDI1
Protein-wise Z-score
Cluster 5
DNMT3B
DNMT3A
PRORP
UHRF1
DDX21
MYBBP1A
SS18
PTBP1
RRP7A
POLR1G
UTP14A
UTP20
POLR1E
UTP15
WDR46
Protein-wise Z-score
Cluster 2
RPS6KA3
DUSP3
MAP2K4
PRDX2
SESN2
PSMD10
NUDT2
MAP2K6
ATOX1
BLVRA
PSMD9
RELA
TRAF6
BIRC2
IDH1
Protein-wise Z-score
Cluster 6
ASNS
ETF1
EIF5B
EEF2
RPL27
RPLP0
RPS3
EEF1D
EIF4B
TRMT112
GCN1
RPLP2
RPL17
RPL12
RPL31
Protein-wise Z-score
Cluster 3
SLC25A4
GOT2
NUBPL
SLC25A22
BCS1L
APOO
SLC25A11
PDP1
LETM1
NDUFAF4
APP
IDH3G
TMEM126A
ATP5MF
NDUFS7
Protein-wise Z-score
Cluster 7
ARID3A
ADA
PGM1
SHMT1
PAICS
RRM1
MPI
E2F4
LDHA
MAT2B
MALT1
PPAT
NUDT5
PHGDH
BCAT1
Protein-wise Z-score
Cluster 4
CYCS
NDUFA4
AKAP1
COX20
MRPS24
TIMMDC1
PAM16
IDH3B
MICU2
NDUFB6
TOMM5
MRRF
PARL
ARG2
MRPL34
Protein-wise Z-score
Cluster 8
H2AX
H1-2
KDM1A
KDM5B
H1-4
H1-5
UBE2I
ARID4B
H3C12
LMNB1
CBX5
H2BC11
HDAC2
ARID1B
ENY2
Protein-wise Z-score

**Supplementary Figure 1: coordinated remodeling of mtDNA content and the proteome during human motor neuron differentiation.** **(A)** mtDNA copy number measured by qPCR in ESC H1–derived motor neurons up to day 16, confirming the trend observed in Figure 1C. $n = 3$ technical replicates (wells). **(B)** mtDNA copy number measured by qPCR at key stages of motor neuron differentiation (d0, d10, and >d30) in iPSC3 and ESC H1. Validation of the results observed in Figure 1C. $n = 3$ technical replicates (wells). **(C)** Effect of 48 hours of 0.5 µM ROCKi treatment on mtDNA copy number in PSCs, as neuronal differentiation could not be induced in the absence of ROCKi. mtDNA copy number was assessed by qPCR. Cell line: iPSC1; 3 technical replicates (wells); analyzed using a *t*-test. **(D)** mtDNA copy number measured by qPCR during after the 2 first days of neuronal differentiation in conditions where CHIR, SB, or LDN were progressively removed. Any combination of removals prevented the drop in mtDNA copy number. Cell line: iPSC1; 2-3 technical replicates (wells). **(E)** mtDNA copy number measured by qPCR until day 4 of neuronal differentiation in conditions where SAG and RA were progressively removed. No effect on mtDNA copy number was observed. Cell line: iPSC1; 3 technical replicates (wells). **(F)** mtDNA copy number measured by qPCR where SAG and RA were either progressively removed until day 14 (left panel) or extended until day 16 (right panel). No effect on mtDNA copy number was observed. Cell line: iPSC1; 3-6 technical replicates (wells). **(G)** Left: representative images of human PSC spheres prior to grafting, day 10 tumors, and week 5 tumors. Right: mtDNA copy number measured by qPCR in undifferentiated PSCs (und) and teratomas at different time points. qPCR primers were specific to human DNA, excluding contamination from mouse DNA. $n = 3$ biological replicates (except at week 6, where $n = 2$ due to one animal loss), including iPSC1, iPSC2, and ESC H1. For **A-G**, error bars represent standard deviation. Statistical significance was defined as $p < 0.05$ (*), $p < 0.01$ (**), or $p < 0.001$ (***), ns = non-significant. Multiple group comparisons were performed using one way ANOVA followed by Dunnet's multiple comparison test (comparisons vs control group), unless otherwise indicated. **(H)** Individual temporal profiles of pathway-associated proteins within each cluster. For each trajectory cluster, the 15 proteins associated with the selected Reactome and KEGG pathways showing the greatest temporal variation were selected. Proteins were ranked according to the range between their highest and lowest mean $\log_2$ abundance across differentiation. Heatmaps show protein-wise Z-scores calculated across five independent wells.

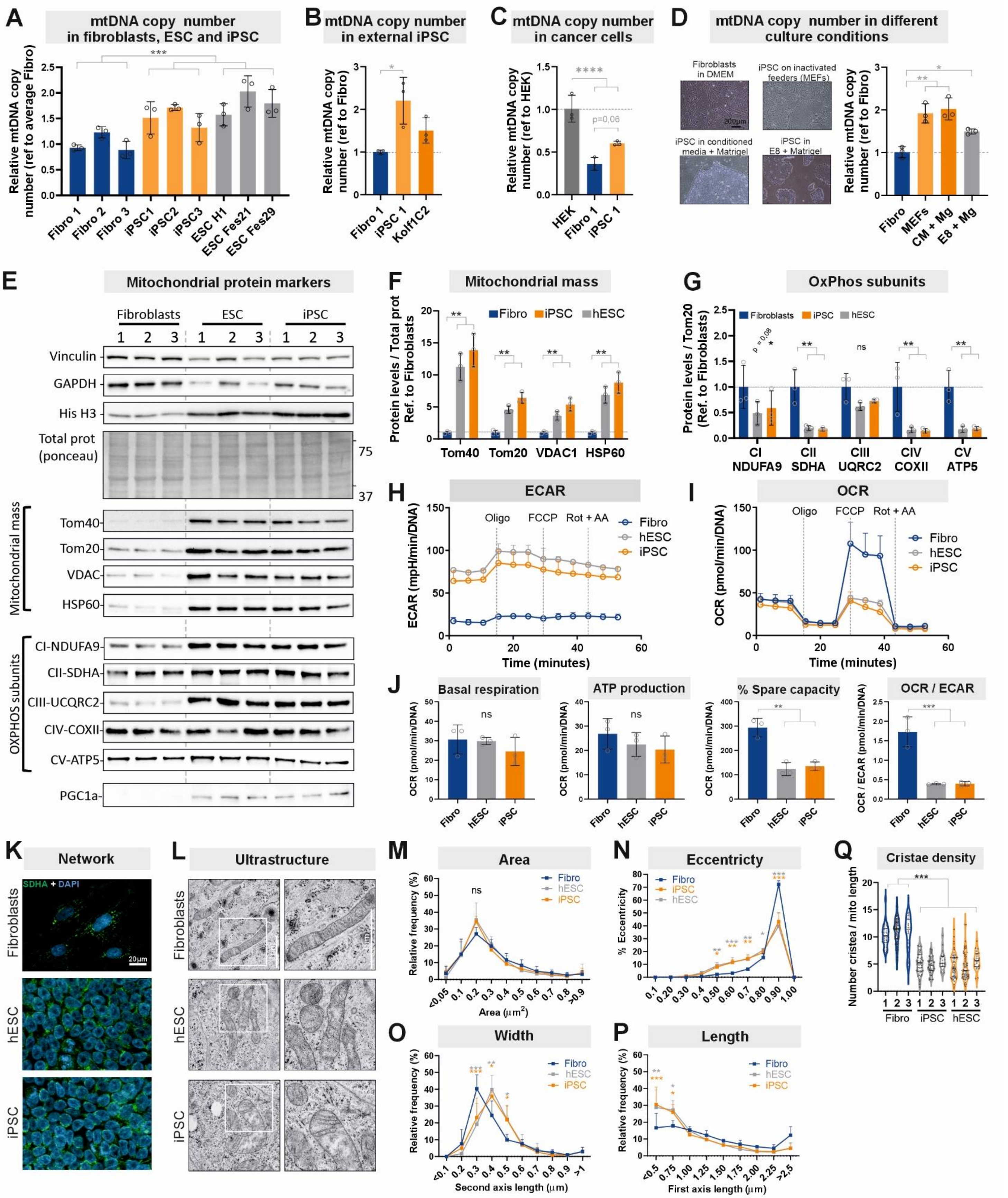
A
mtDNA copy number in fibroblasts, ESC and iPSC
Relative mtDNA copy number (ref to average Fibro)
Fibro 1
Fibro 2
Fibro 3
iPSC1
iPSC2
iPSC3
ESC H1
ESC Fes21
ESC Fes29
B
mtDNA copy number in external iPSC
Relative mtDNA copy number (ref to Fibro)
Fibro 1
iPSC 1
Kolf1C2
C
mtDNA copy number in cancer cells
Relative mtDNA copy number (ref to HEK)
p=0,06
HEK
Fibro 1
iPSC 1
D
mtDNA copy number in different culture conditions
Fibroblasts in DMEM
iPSC on inactivated feeders (MEFs)
iPSC in conditioned media + Matrigel
iPSC in E8 + Matrigel
200μm
Relative mtDNA copy number (ref to Fibro)
Fibro
MEFs
CM + Mg
E8 + Mg
E
Mitochondrial protein markers
Fibroblasts
ESC
iPSC
Vinculin
GAPDH
His H3
Total prot (ponceau)
75
37
Mitochondrial mass
Tom40
Tom20
VDAC
HSP60
OXPHOS subunits
CI-NDUFA9
CII-SDHA
CIII-UCQRC2
CIV-COXII
CV-ATP5
PGC1a
F
Mitochondrial mass
Fibro
iPSC
hESC
Protein levels / Total prot Ref. to Fibroblasts
Tom40
Tom20
VDAC1
HSP60
G
OxPhos subunits
Fibroblasts
iPSC
hESC
Protein levels / Tom20 (Ref. to Fibroblasts)
p = 0,08
ns
CI NDUFA9
CII SDHA
CIII UQRC2
CIV COXII
CV ATP5
H
ECAR
ECAR (mpH/min/DNA)
Oligo
FCCP
Rot + AA
Fibro
hESC
iPSC
Time (minutes)
I
OCR
OCR (pmol/min/DNA)
Oligo
FCCP
Rot + AA
Fibro
hESC
iPSC
Time (minutes)
J
Basal respiration
ATP production
% Spare capacity
OCR / ECAR
OCR (pmol/min/DNA)
OCR / ECAR (pmol/min/DNA)
ns
Fibro
hESC
iPSC
K
Network
SDHA + DAPI
Fibroblasts
hESC
iPSC
20μm
L
Ultrastructure
Fibroblasts
hESC
iPSC
M
Area
ns
Fibro
hESC
iPSC
Relative frequency (%)
Area (μm2)
N
Eccentricity
Fibro
iPSC
hESC
% Eccentricity
O
Width
Fibro
hESC
iPSC
Relative frequency (%)
Second axis length (μm)
P
Length
Fibro
hESC
iPSC
Relative frequency (%)
First axis length (μm)
Q
Cristae density
Number cristea / mito length
Fibro
iPSC
hESC

**Supplementary Figure 2: pluripotent stem cells have a large number of mitochondria but it is immature. (A)** mtDNA copy number measured by qPCR in three iPSC lines and their corresponding parental somatic fibroblasts, as well as in three ESC lines. Data represent n = 3 technical replicates (wells). Data were normalized to the average of the three fibroblast cell lines. **(B)** Left: representative bright-field images of fibroblast and iPSC cultured under different conditions. Scale bar = 200 µm. Right: mtDNA copy number quantified by qPCR in iPSC cultured on inactivated feeder mouse embryonic fibroblasts (MEFs) or under feeder-free conditions using either MEF-conditioned medium with Matrigel (CM + Mg) or Essential 8 medium with Matrigel (E8 + Mg). Data represent n = 3 biological replicates (iPSC1-3), each calculated as the average of three technical replicates (wells). **(C)** mtDNA copy number measured by qPCR in Fibro 1, reprogrammed iPSC1 and an external iPSC line, Kolf1C2. Data represent n = 3 technical replicates (wells). We observed similar results in an externally derived and curated iPSC line (Kolf1C2), confirming that this was not specific to our in-house lines. **(D)** mtDNA copy number measured by qPCR in HEK239 cancer cells, Fibro 1 and reprogrammed iPSC1. Data represent n = 3 technical replicates (wells). The mitochondrial content of PSCs remained lower than that of the cancer cell line HEK293, indicating that the increase in PSCs is still modest compared to highly proliferative transformed cells. **(E)** Western blot analysis was performed on the same cell lines used in Sup. Figure 1A (n = 3 biological replicates). Common housekeeping proteins were not reliable; therefore, total protein normalization based on Ponceau staining was used. Several mitochondrial mass markers were analyzed, including proteins localized to the outer mitochondrial membrane (Tom40, Tom20, and VDAC) and the mitochondrial matrix (HSP60). In addition, one representative subunit from each OxPhos complex was assessed (CI = complex I, and so forth). PGC1α was included as a master regulator of mitochondrial biogenesis. **(F)** Quantification of the Western blot analysis normalized to total protein (Ponceau staining), as a measure of mitochondrial mass per cell. n = three independent experiments, with each value calculated as the mean of three biological replicates (three cell lines). Similar to mtDNA content, mitochondrial mass markers were elevated in PSCs. **(G)** Quantification of the Western blot analysis, normalized to mitochondrial mass (Tom20), as a measure of OxPhos subunits per mitochondrion. n = three independent experiments, with each value calculated as the mean of three biological replicates (three cell lines). Interestingly, although PSCs have increased mitochondrial abundance, the overall levels of OxPhos subunits are reduced. **(H)** Extracellular acidification rate (ECAR) kinetics measured using the Seahorse XF Analyzer. **(I)** Oxygen consumption rate (OCR) kinetics measured using the Seahorse XF Analyzer. **(J)** OCR- and ECAR-derived parameters showing comparable basal respiration across cell types, but a reduced % spare respiratory capacity (stress-induced increase in respiration) in PSCs, along with a decreased OCR/ECAR ratio, indicating that PSCs primarily rely on glycolysis. For **H-J**, n = 3 three biological replicates (same as for Sup. Figure 1A). Each individual dot represents the combined average of 6-12 technical replicates (wells) per cell line. **(K)** Representative fluorescence images showing the mitochondrial network in fibroblasts (#1), iPSC (#1) and ESC (H1). SDHA, a Complex II subunit, is used as a marker of mitochondrial mass. DAPI labels nuclei. Scale bar = 20 μm. Mitochondrial network looks more complex and elongated in fibroblasts. **(L)** Representative transmission electron microscopy images of fibroblasts and PSCs, including magnified views. Scale bar = 1 μm. **(M–P)** Quantification of mitochondrial morphology parameters, including area, first axis (length), second axis (width), and eccentricity (where 0 indicates a perfect line and 1 indicates a perfect circle). Although the average area remains comparable, PSC contain more rounded and shorter mitochondria compared to fibroblasts. For (M–P), $n$ = 3 biological replicates (same cell lines as in Sup. Fig. 1A), as the average of 840–1,991 mitochondria analyzed per cell type; showing reduced cristae density in PSCs compared to fibroblasts. **(Q)** Number of cristae per mitochondrial length. 20–81 mitochondria were analyzed per cell line. For all, error bars represent standard deviation. Statistical significance was defined as $p < 0.05$ (*), $p < 0.01$ (**), or $p < 0.001$ (***), ns = non-significant. Multiple group comparisons were performed using one-way ANOVA followed by Dunnet´s multiple comparison test (comparisons vs control group) or Tukey's multiple comparison test (comparison all groups vs all groups).

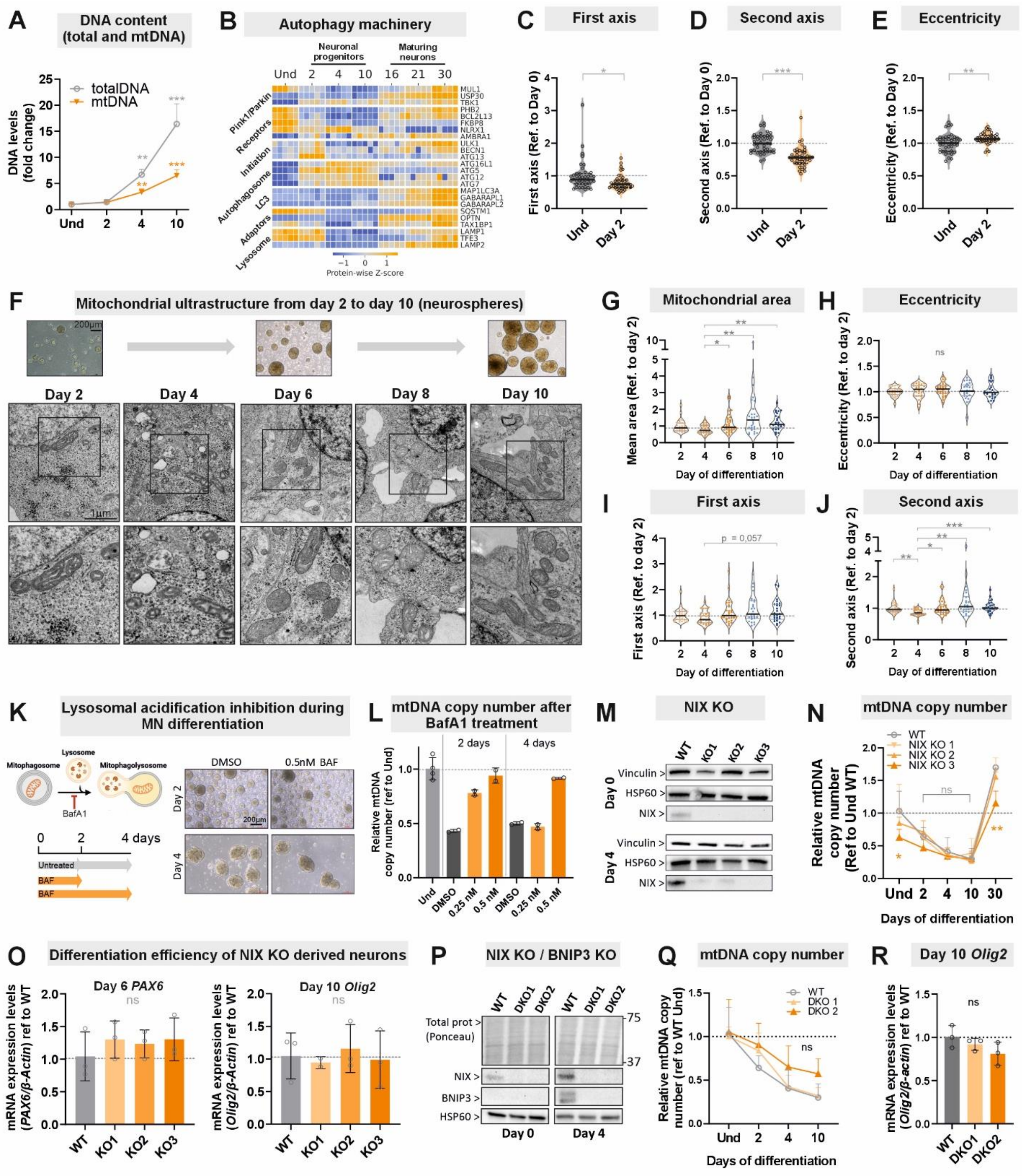


**Supplementary Figure 3: mitochondrial depletion during the exit of pluripotency is driven by fragmentation and lysosomal acidification: (A)** Total DNA content was quantified using the CyQuant fluorescent dye kit, and absolute mtDNA content was measured by qPCR (not normalized to nuclear DNA). Data are presented as fold change during the first 10 days of differentiation. n = 3 technical replicates (wells) of iPSC1. **(B)** Heatmap of proteomics data showing detected autophagy-related factors, displayed as Z-scores. Proteins were selected from the canonical PINK1/Parkin-dependent and receptor-mediated mitophagy pathways, together with core autophagy initiation, autophagosome formation, cargo recognition, and lysosomal machinery components, based on current literature and pathway annotations (Jiang et al., 2021, Yoshii and Mizushima, 2017). **(C-E)** Quantification of mitochondrial morphology parameters between undifferentiated iPSC1 (day 0) and day 2 neural progenitors cultured as a monolayer; including first axis (length), second axis (width), and eccentricity (where 0 indicates a perfect line and 1 indicates a perfect circle). Extension of Figure 2I-J. n = 42 images containing 439 (day 0) or 272 (day 2) mitochondria from iPSC1 derived neurons. Differences between time points were assessed using an unpaired Welch's t-test on log-transformed image-level means. **(F)** Representative transmission electron microscopy images of iPSC1-derived neurons from day 2 until day 10 growing as neurospheres in suspension, including magnified views. Scale bar = 1 μm. **(G-J)** Quantification of mitochondrial morphology parameters, including area, first axis (length), second axis (width), and eccentricity (where 0 indicates a perfect line and 1 indicates a perfect circle), extracted from (H). n = 26-30 images containing ~150 mitochondria in total per stage from iPSC1 derived neurons. Statistical analyses were performed on $log_2$-transformed image-level values using Brown–Forsythe and Welch one-way ANOVA followed by Dunnett's T3 multiple-comparison test. **(K)** Left, cartoon depicting the experimental setup to inhibit lysosomal acidification with Bafilomycin A1 (BafA1). Created in BioRender. Torregrosa, R. (2026) https://BioRender.com/rnmmctn. Right, representative bright-field images of BafA1-treated cells after 2 or 4 days of MN differentiation. The morphology and overall health of neurospheres were not affected by BAF treatment. Scale bar = 200 μm**. (L)** mtDNA copy number determined by qPCR in iPSC-derived neurons after 2 or 4 days of 0.25 or 0.5 μM BAF treatment. Cell line: iPSC1; n = 2-4 technical replicates (wells). **(M)** Western blot analysis of day 0 (undifferentiated iPSC) and day 4 neuronal progenitors validating the knockout of NIX (BNIP3L). WT represents the parental iPSC1 line and KO1–3 are three independent clones in which NIX expression was dramatically reduced (KO 1 and 2) effectively blunted (KO3). **(N)** mtDNA copy number measured by qPCR in WT (parental) and the three NIX KO cell lines. Loss of NIX did not alter mtDNA content dynamics during differentiation. n = 2-3 technical replicates (wells). All samples referred and compared to WT Und. **(O)** mRNA expression levels in WT (parental) and the three NIX KO cell lines of the early neuroectoderm marker *PAX6* (day 6) and the neuronal progenitor marker *OLIG2* (day 10). **(P)** Western blot analysis of day 0 (undifferentiated iPSC) and day 4 neuronal progenitors validating the double knockout (DKO) of NIX (BNIP3L) and BNIP3. BNIP3 was knocked out in the NIX KO 3 cell line. WT represents the parental iPSC1 line and DKO1–2 are two independent clones in which NIX and BNIP3 expression was blunted. **(Q)** mtDNA copy number measured by qPCR in WT (parental) and the two DKO cell lines. Loss of NIX and BNIP3 did not alter mtDNA content dynamics during differentiation. n = 3 technical replicates (wells). All samples referred and compared to WT Und. **(R)** mRNA expression levels in WT (parental) and the two DKO cell lines of the neuronal progenitor marker *OLIG2* (day 10). Data represent $n = 3$ technical replicates (wells). In all cases, error bars represent standard deviation. Statistical significance was defined as $p < 0.05$ (*), $p < 0.01$ (**), or $p < 0.001$ (***), ns = non-significant. Multiple group comparisons were performed using one-way ANOVA followed by Dunnet´s multiple comparison test (comparisons vs control group), unless otherwise indicated.

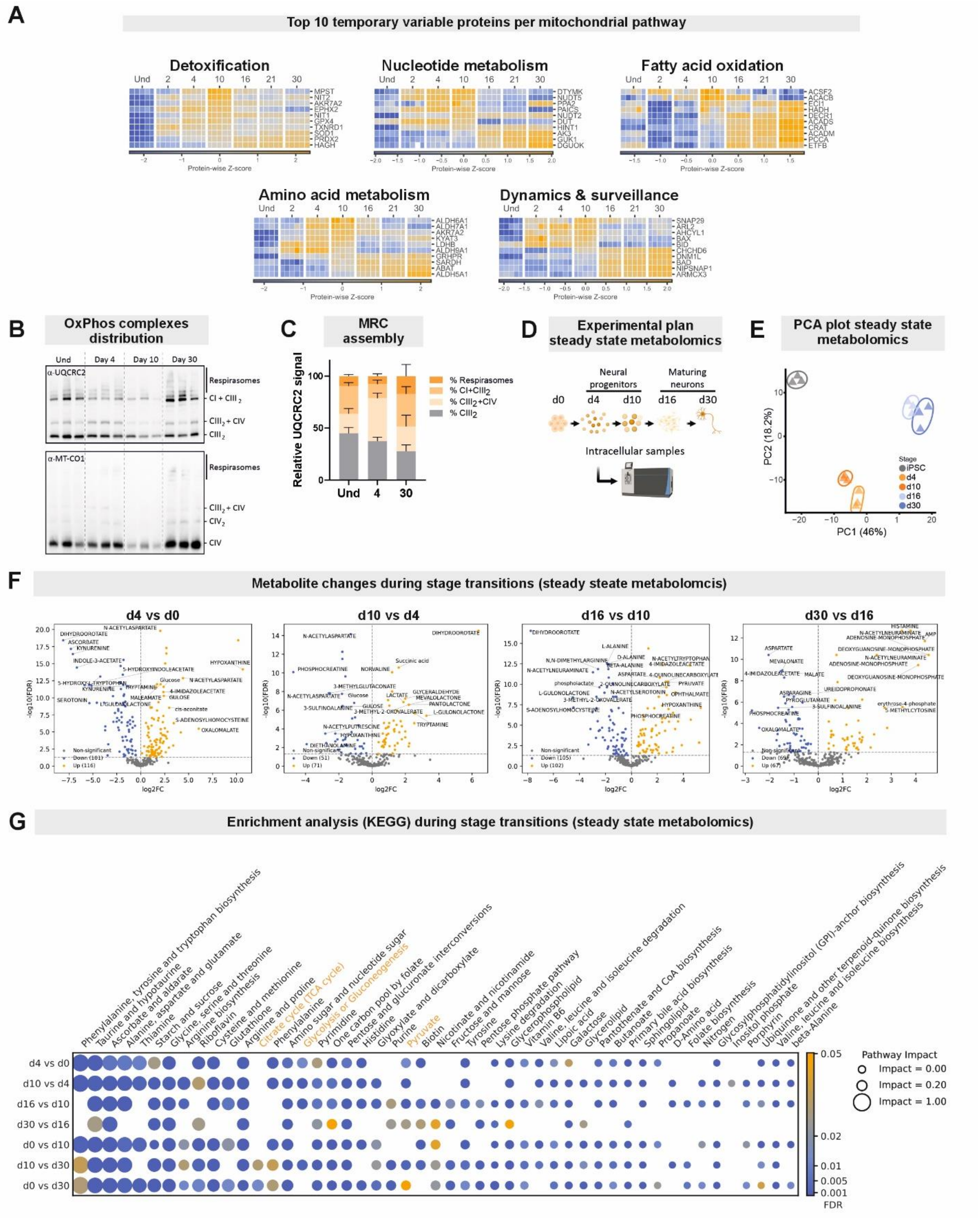


**Supplementary Figure 4: Coordinated remodeling of mitochondrial OxPhos complexes during neuronal maturation. (A)** MitoCarta 3.0 MitoPathway annotations were collapsed into broad functional modules using a predefined hierarchical classification (see Figure 3E and F). Heatmaps show proteins from selected modules: Detoxification, Nucleotide metabolism,

Fatty acid oxidation, Amino acid metabolism, Central dogma, and Dynamics & surveillance. For each module, proteins were ranked by the change in mean log2 abundance at D10 relative to undifferentiated iPSC, and the top 10 proteins are shown. Protein abundance was displayed as Z-scores. Columns represent individual biological replicates (n = 5 per time point). **(B)** Representative Blue-Native PAGE (BN-PAGE) immunoblot showing respiratory chain complexes organization. UQCRC2 is shown as a marker of Complex III–containing assemblies, while MT-CO1 is used as a marker of CIV-containing assemblies. n = 3 biological replicates (iPSC1-3). **(C)** Mitochondrial respiratory chain (MRC) assembly. Quantification of the BN-PAGE analysis shown in (A) using the UQCRC2 blot, depicting the relative abundance (%) of respirasomes ($CI+CIII_2+CIV$), supercomplexes $I+III_2$ and $III_2+IV$, and free $CIII_2$. **(D)** Steady state metabolomics experimental design showing the selected time points. Created in BioRender. Torregrosa, R. (2026) https://BioRender.com/qwqb483. Cell line: iPSC1; 3 technical replicates (wells). **(E)** Principal component analysis (PCA) of the metabolomics data showing clear separation between differentiation stages. **(F)** Volcano plots showing differential metabolite abundance between consecutive stages of motor neuron differentiation. Differential abundance was determined using limma with empirical Bayes moderation and Benjamini–Hochberg correction. Axes show $\log_2$ fold change and $-\log_{10}$(FDR). Vertical dashed line indicates the fold-change threshold ($|\log_2 FC| \geq 1$), and the horizontal dashed line indicates the significance threshold (FDR < 0.05). The most significant annotated metabolites are labelled. **(G)** Differential metabolites were subjected to KEGG pathway analysis using MetaboAnalyst.com (Pang et al., 2024). Pathway enrichment was performed using over-representation analysis (hypergeometric test), and pathway topology was evaluated using relative betweenness centrality. P values were adjusted using the Benjamini–Hochberg method, and pathway impact scores were derived from the topology analysis.

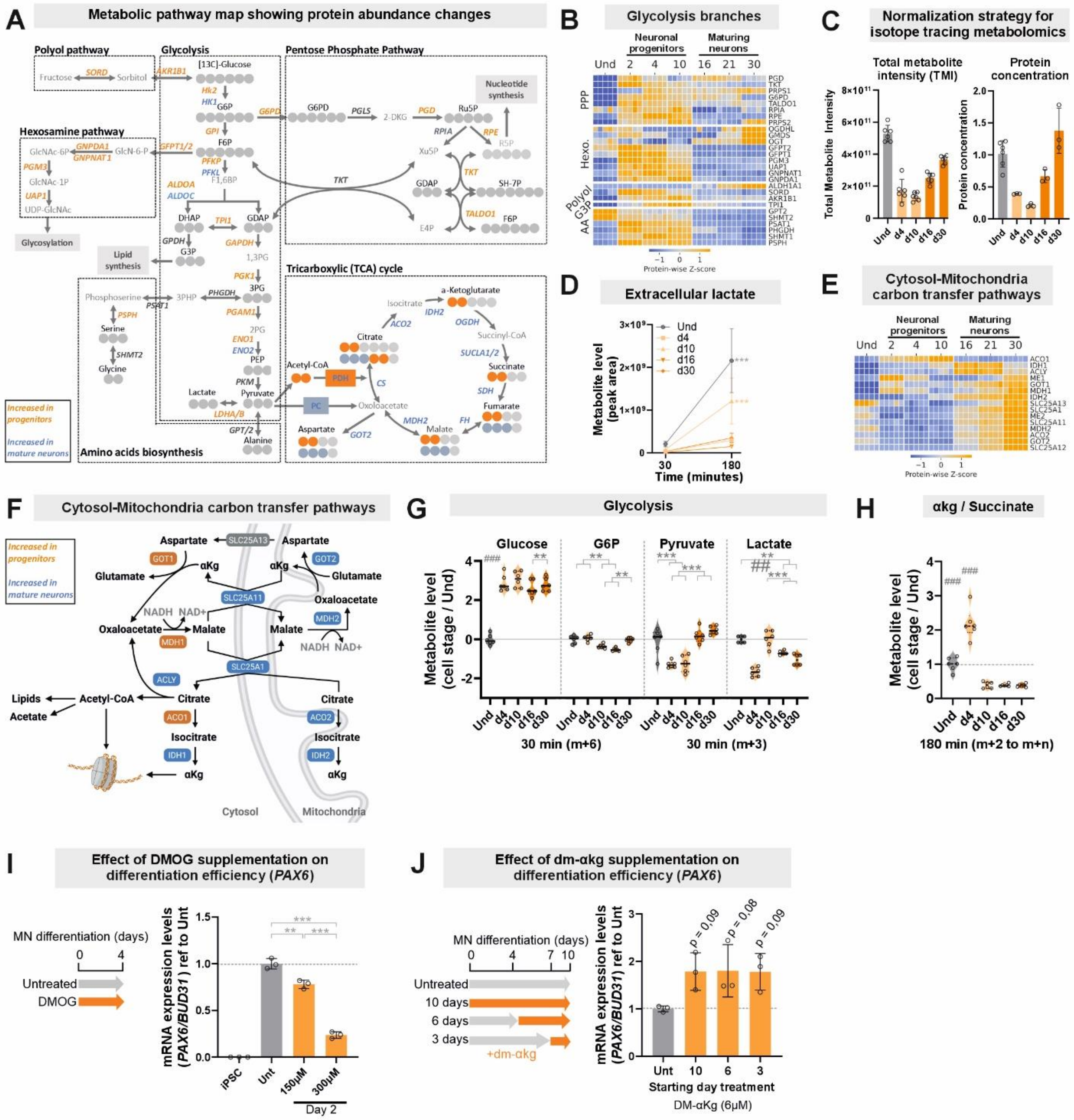


**Supplementary Figure 5: Metabolic shift from glycolysis-dominant early progenitors to enhanced OxPhos during neuronal maturation.**

**A)** Cartoon schematic of glycolysis, glycolytic branches and TCA cycle metabolites and enzymes. Shadowed circles denote carbon atoms as in Figure 4G. Protein names labelled in orange are increased in the neural progenitor stages, whereas blue denotes increased in the more mature stages. **B)** Heatmap from proteomics data showing glycolysis branches enzymes of the pentose phosphate pathway (PPP), hexosamine pathway, sorbitol pathway (Sor.)

glycerol-3-phosphate pathway (G3P), and amino acid synthesis (AA), displayed as Z-scores. Columns represent individual biological replicates (n = 5 per time point). **C)** TMI and protein concentration measurements of metabolomics samples used for steady-state and isotope-tracing analyses. TMI and protein concentration followed broadly similar temporal patterns, indicating that changes in metabolite abundance largely reflected differences in cellular biomass. However, the magnitude of these changes varied substantially across time points. Therefore, quantile normalization was applied for downstream statistical analyses. **D)** Extracellular levels of lactate after 30 or 180 minutes in medium containing $U^{13}C$-glucose. Metabolites levels normalized to their corresponding 30 minutes values (peak area). **E)** Heatmap from proteomics data showing selected proteins involved in cytosol-mitochondria carbon transfer pathways, including the malate-aspartate and citrate-malate shuttles, displayed as Z-scores. Columns represent individual biological replicates (n = 5 per time point). **F)** Cartoon featuring the enzymes and metabolites involved in cytosol-mitochondria carbon transfer pathways. Created in BioRender. Torregrosa, R. (2026) https://BioRender.com/3vu5v8h. Protein names labelled in orange are increased in the neural progenitor stages, whereas blue denotes increased in the more mature stages. Proteins labelled in grey were not detected in the data set. **G)** Relative abundance of relevant $^{13}C$-labeled isotopologues (m+6 or m+3) in glycolysis metabolites, shown as log2-transformed values normalized to undifferentiated (Und) cells. **H)** Ratios of labelled (m+2 to m+n) α-ketoglutarate (αKg) to succinate extracted from the $^{13}C$ isotope tracing experiment. Lower ratios indicate increased αKg consumption via the 2-oxoglutarate dehydrogenase (OGDH) reaction or reductive carboxylation, which alters the αKG/succinate balance to potentially modulate αKG-dependent dioxygenases and drive epigenetic changes such as histone or DNA demethylation. **I)** Effect of DMOG, an inhibitor of αKG-dependent dioxygenases, on neuronal differentiation efficiency based on (TeSlaa et al., 2016). Left: schematic of the experimental design. DMOG

(150–300 μM) was added from day 0 to day 4 of the differentiation protocol, coinciding with the transient peak in αKG levels, followed by qPCR quantification of PAX6 mRNA expression. Cell line: iPSC1; $n = 3$ technical replicates (wells). **J)** Effect of dimethyl-αKG (dm-αKG) supplementation on neuronal differentiation efficiency. Left: schematic of experimental setup. 6 μM dm-αKG were added for 3, 6, or 10 days up to day 10 of the differentiation protocol, followed by qPCR quantification of *PAX6* mRNA levels. Cell line: iPSC1; $n = 3$ technical replicates (wells). Error bars represent standard deviation. Statistical significance was defined as $p < 0.05$ (*), $p < 0.01$ (**), or $p < 0.001$ (***); # denotes significantly different compared to all groups. Multiple group comparisons were performed using one way ANOVA followed by Tukey's multiple comparison test.

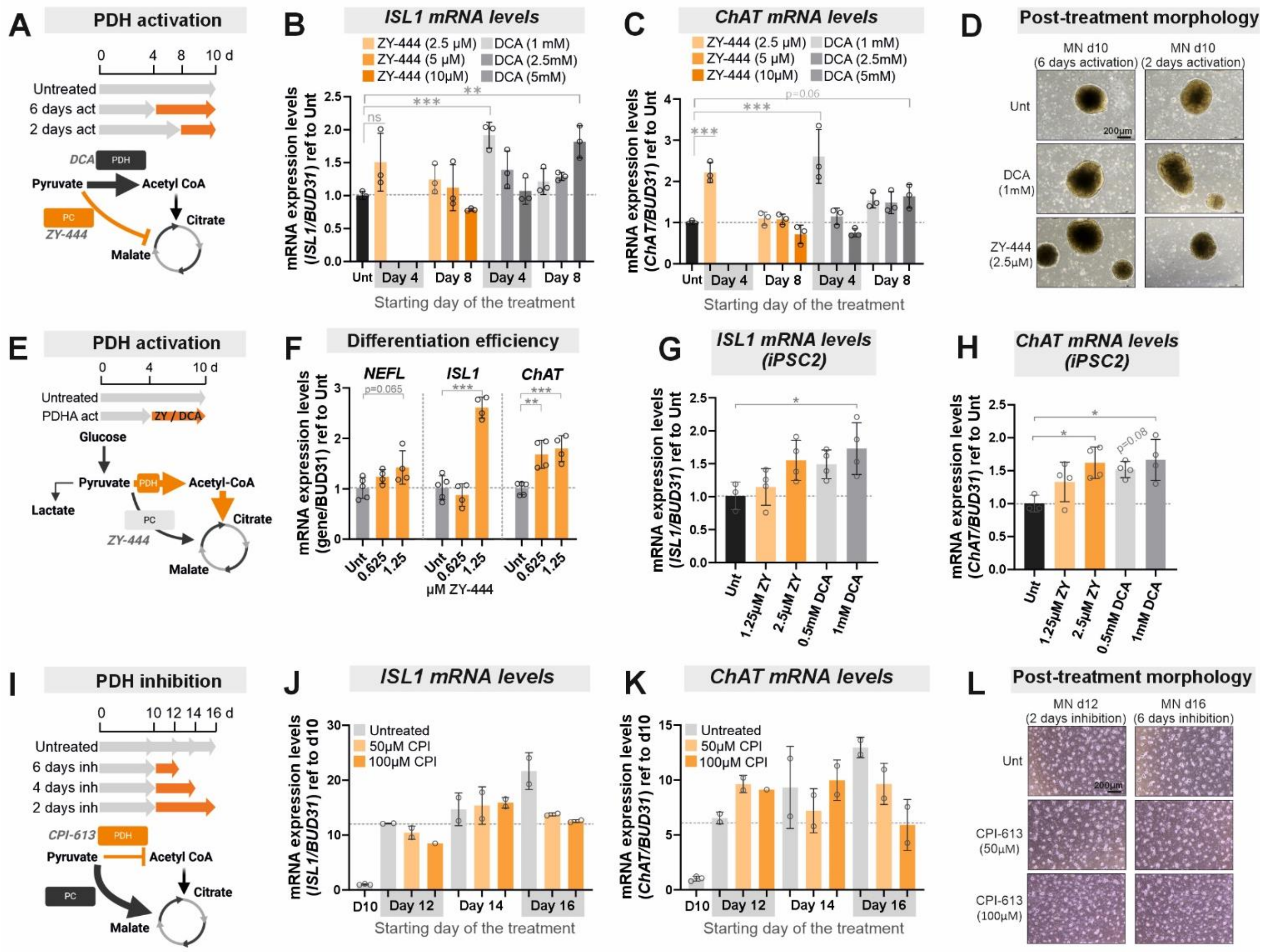


**Supplementary Figure 6: Mitochondrial pyruvate metabolism influences the pace of neuronal maturation. A)** Cartoon schematic of the titration of PDH activation during the progenitor stage. Created in BioRender. Torregrosa, R. (2026) https://BioRender.com/jqyagdk. Dichloroacetate (DCA) or ZY-444 were added to the culture medium at different concentrations and time windows (from day 4 or day 8 to day 10), followed by gene expression analysis by qPCR from day 10 samples. DCA activates PDH by inhibiting PDKs (Mathieu and Ruohola-Baker, 2017, Stacpoole et al., 2008, TeSlaa et al., 2016), whereas ZY-444 directly inhibits PC (Lin et al., 2020, Liu et al., 2022), reducing pyruvate-to-acetyl-CoA flux, potentially increasing reliance on alternative anaplerotic pathways such as pyruvate carboxylase. **B–C)**

Differentiation efficiency following treatment (as described in A) was assessed by qPCR analysis of neuronal maturation markers (ISL1 and ChAT). High concentrations of ZY-444 for the longer period was toxic. Cell line: iPSC1; n = 3 technical replicates (wells). **D)** Representative bright-field images acquired post-treatment revealed no clear changes in sphere morphology or cell death size at the concentrations used in subsequent experiments. **E)** Cartoon schematic of the PC inactivation strategy during the progenitor stages. Created in BioRender. Torregrosa, R. (2026) https://BioRender.com/jqyagdk. ZY-444 was added to the culture medium from day 4 to day 10. **F)** Differentiation efficiency following PC inhibition, assessed by qPCR analysis of the expression levels of neuronal maturation markers (NEFL, ISL1, and ChAT) on day 10. Cell line: iPSC1; n = 3 technical replicates (wells). **G-H)** Validation of the effect of ZY-444 (ZY) and dichloroacetate (DCA) on accelerating neuronal maturation in an independent cell line (iPSC2, n= 3 technical replicates). Cells were treated for 6 days following the same experimental setup described in Figure 5A and using the indicated concentrations. Neuronal maturation was assessed on day 10 by measuring the expression levels of *ISL1* and *ChAT*. **I)** Cartoon schematic of the titration of PDH inhibition during the maturation stage. Created in BioRender. Torregrosa, R. (2026) https://BioRender.com/jqyagdk. CPI-613 was added to the culture medium at different concentrations and time windows (from day 10 until day 12, 14 or 16), followed by gene expression analysis by qPCR on days 12, 14 or 16. CPI-613 is a lipoate analogue that inhibits mitochondrial metabolism by targeting the pyruvate dehydrogenase (PDH) (and α-ketoglutarate dehydrogenase (KGDH)), thereby reducing pyruvate-to-acetyl-CoA flux, potentially increasing reliance on alternative anaplerotic pathways such as pyruvate carboxylase (Elia et al., 2022, Zachar et al., 2011). **J-K)** Differentiation efficiency following CPI-613 treatment was assessed by qPCR analysis of neuronal maturation markers (ISL1 and ChAT). Cell line: iPSC1; n = 3 technical replicates (wells). With n=2 the statistical analysis was not applied, but 6 days treatment with 100 µM

CPI-613 clearly reduced gene marker expression levels. **L)** Representative bright-field images acquired post-treatment revealed no clear changes in the neuronal cultures at the concentrations used in subsequent experiments (100 μM CPI-613). Data are presented as mean ± standard deviation and normalized to untreated (Unt) cells. Statistical significance was defined as $p < 0.05$ (*), $p < 0.01$ (**), or $p < 0.001$ (***). Pairwise comparisons were performed using a t-test. Multiple group comparisons were conducted using one-way ANOVA followed by Dunnett's multiple comparison test against untreated control.

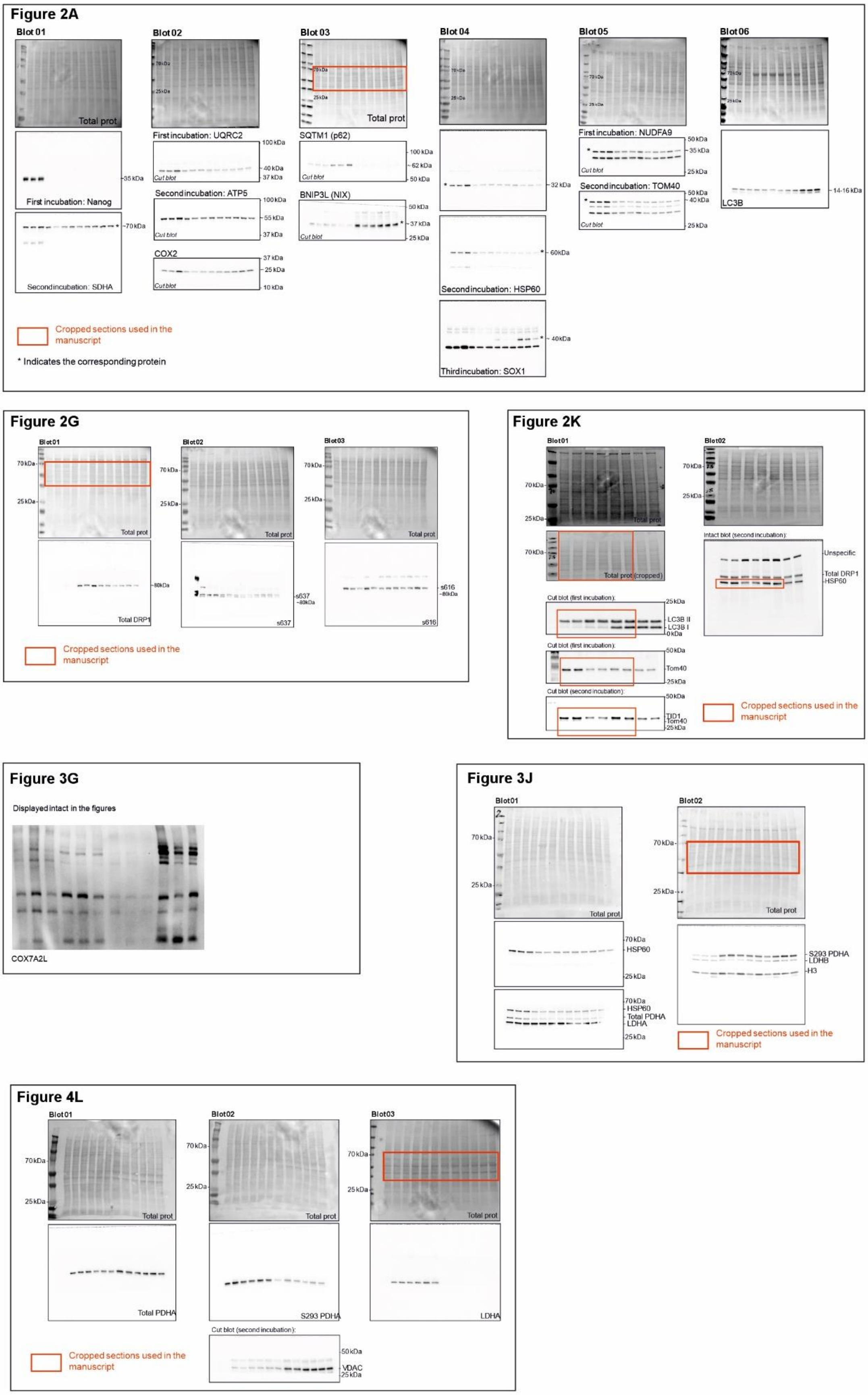


**Supplementary Figure 7: Uncropped Western blot images corresponding to the main figures**

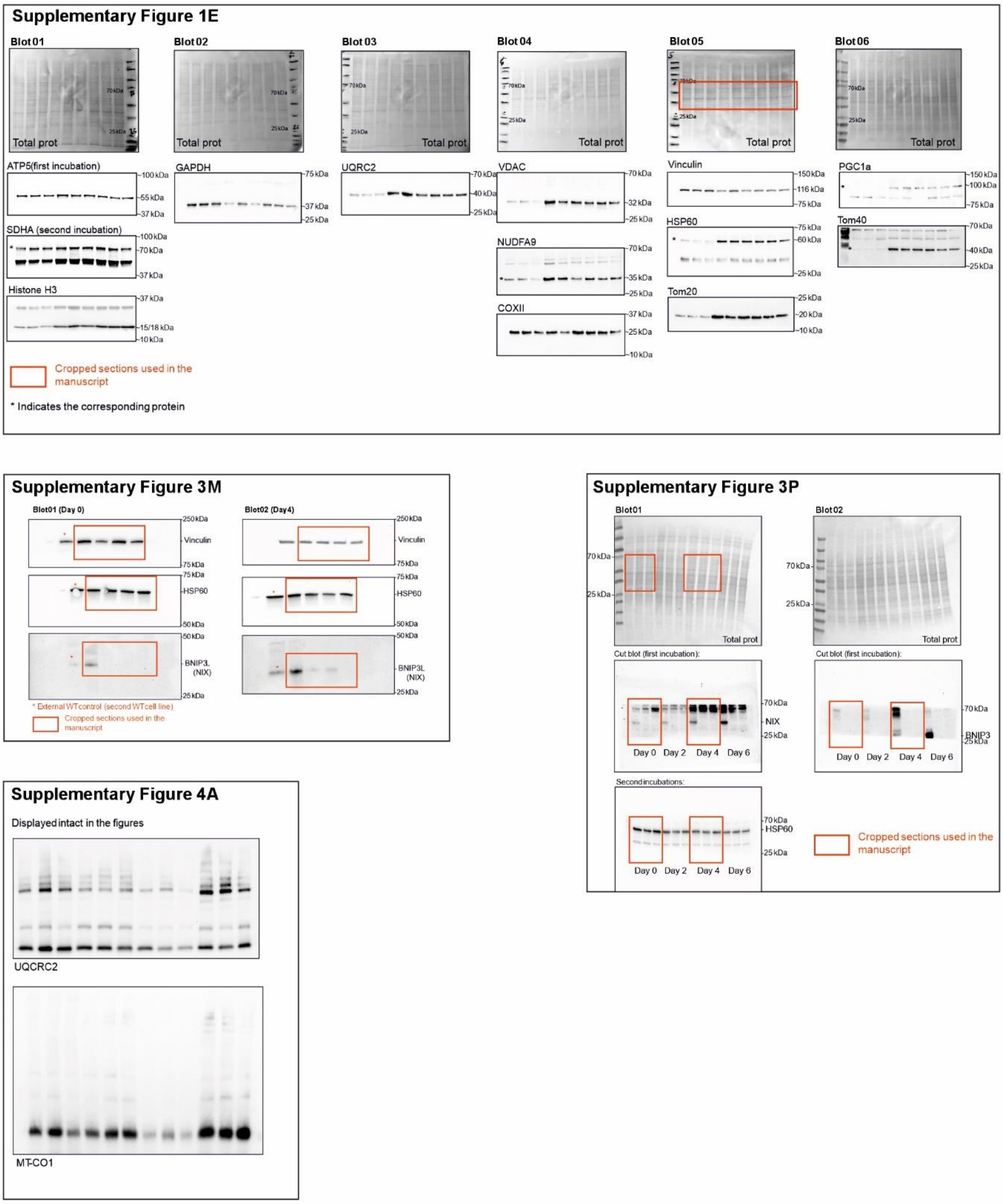


**Supplementary Figure 8: Uncropped Western blot images corresponding to the supplementary figures.**
